# Development of Dendritic Cell Membrane-Coated Nanoparticles for Antigen-Specific T Cell Engagement

**DOI:** 10.1101/2025.08.06.664969

**Authors:** Sao Puth, Shruti Sunil Jadhav, Ali Zareein, Jimmy Blauser-Wilson, Mina Mahmoudi, Ruben Rojas Betanzos, Bayonel Ventura, Andrea M. Sprague-Getsy, Xiaoran Hu, James L. Hougland, Yaoying Wu

## Abstract

Dendritic cell (DC) membrane-coated nanoparticles (DCmPs) hold significant potential for antigen-specific therapies. DCmPs carry key DC membrane proteins that facilitate DC-T cell interaction, such as major histocompatibility complex (MHC), costimulatory CD80/86, and adhesive molecules ICAM-1. However, our current understanding of the impact of the coating processes and the composition of the final products is very limited, significantly hindering the development of DCmP-based therapy. Here, using DC2.4 cell membrane proteins and poly (lactic-co-glycolic acid) (PLGA) nanoparticles, we comprehensively characterized and compared the compositions and functions of DCmPs produced using sonication, extrusion, and a newly developed combined coating approach (sonication coating following by extrusion process). The combined coating approach achieved relatively high level of protein coating and exerted superior control over the diameter and uniformity of DCmPs relative to sonication and extrusion. We also developed a characterization strategy by leveraging the homotypic interactions between DCmPs and DC2.4 cells and determined that about 80% of PLGA particles are coated with membrane proteins and both unbound proteins and uncoated particles similarly present in the final products after the three coating processes. Because DC2.4 cells exclusively express MHC class I molecules, DCmPs showed preferentially bind to cognate B3Z CD8+ T cells over DOBW CD4+ T cells, confirming that DCmPs bind to T cells in an antigen-specific fashion. Furthermore, we demonstrated that DCmPs can activate B3Z CD8+ T cells *in vitro*, similar to DC2.4 cells. These findings demonstrate a new coating approach that potentially improves size control over membrane-coated particles and a characterization strategy for detailed analysis of coated particle composition, having important and broad implications for the therapeutic development of DCmPs and other membrane-coated particle technology.

## Introduction

Cell membrane-coated nanoparticles (CNPs) have emerged as a promising technology for therapeutic applications including drug delivery, cancer immunotherapy, anti-inflammation, and vaccinations.^1–4^ These membrane-coated particles are often composed of synthetic nanoparticulate cores and cell membrane coating. A wide range of particles, including poly(lactic-co-glycolic acid) (PLGA) nanoparticles (NPs), gold NPs, and mesoporous silica NPs, have been explored as the particulate core to facilitate the encapsulation of pharmaceutical agents.^3,5–8^ The cell membrane coating provides CNPs with a variety of membrane proteins and thus the source-cell-mimicking functions. For instance, red blood cell (RBC) membrane-coated NPs benefit from the lack of immune recognition of RBCs and thus prolong their circulation upon systemic administration;^9^ particles coated with tumor cell membrane preferentially accumulate within the source tumor tissues owing to the homotypic membrane interaction and facilitate tumor-targeted drug delivery or tissue imaging;^2,3,10^ macrophage or neutrophil membrane-coated particles can deplete cytokines within local microenvironment and bind to circulating tumor cells owing to the membrane-bound cytokine receptors and the adhesive molecules.^11–16^ Through membrane modification via gene editing, lipid insertion, or membrane hybridization, the functionality of CNPs can be expanded beyond the inherent capabilities of source cells and improve the therapeutic functions or flexibility of CNPs.^17–20^

Recently, dendritic cell (DC) membrane-coated NPs (DCmPs) have drawn growing research attention.^8,21–23^ DCs initiate adaptive immune responses primarily by processing and presenting antigens to T cells using peptide major histocompatibility complex (pMHC).^24^ During this antigen presentation process, in addition to the central interaction between pMHCs and T-cell receptors (TCRs), DCs also provide costimulatory molecule interactions to T cells, such as the binding between CD80/CD86 (DCs) and CD28 (T cells). The DC-T cell interaction is additionally stabilized via adhesive molecule binding between ICAM-1 (Intercellular Adhesion Molecule-1 of DCs) and LFA-1 (Lymphocyte Function-Associated Antigen-1 of T cells).^25,26^ DCmPs can retain these key membrane proteins, including pMHC, and activate antigen-specific T cells for vaccination and immunotherapy.^22,23,27,28^ Additionally, similar to other NP platform, membrane-coated NPs can be delivered via a variety of administration routes, thus enable tissue targeting.^23,29,30^ Collectively, DCmPs represent a new strategy for antigen-specific immunotherapy owing to these advantages. Yet, detailed analyses of the DCmPs regarding the protein coverage and the purity are still lacking, which potentially confounds the interpretation of the observed therapeutic effects and complicates future development.

Membrane-coated particles are commonly produced by mechanically driving the association between particle cores and isolated membrane proteins through sonication or extrusion.^4^ Although the mechanisms underlying the interaction between membrane proteins and particle cores have yet to be fully elucidated, experimental evidence suggests that sonication disrupts membrane structures and promotes protein absorption onto nanoparticles.^3^ Extrusion produce membrane-coated particles by forcing the mixed suspension of proteins and nanoparticles through membranes with hundred-nanometer-sized pores to kinetically trap proteins onto particle cores, which often yields better control over the size of membrane-coated particles.^3,31^ However, comprehensive studies are needed to assess how these two coating approaches influence the particle sizes and the coated protein amount or function. Additionally, the final membrane coating can be influenced by both the cell membrane intrinsic properties, such as the rigidity,^32,33^ and the physical properties of particle cores, including surface charge or elasticity,^5,34^ making it difficult to predict the coating outcome for different coating strategies. Therefore, it is necessary to compare the two coating approaches and devise strategy to achieve consistent coating outcome.

Motivated by the therapeutic potential of DCmPs, we sought to systemically compare the membrane coating process of various coating approaches and study the DCmP functions *in vitro* using a model DCmP produced by coating the cellular membrane of DC2.4 cells, a murine DC cell line, onto PLGA nanoparticles. We developed a combined coating approach by performing additional extrusion steps after sonication, and determined the diameter, coated protein amount, and the purity of final DCmPs from the three coating approaches, i.e. sonication, extrusion, and a combined coating approach. We hypothesize that the combined coating approach can maximize the coated protein amount on DCmPs and create a uniform size profile. We show that DCmPs preserve the key proteins involved in DC-T cell immune synapses formation and preferentially bind to antigen-specific T cells, likely mediated by MHC molecules. Finally, we confirm that DCmPs acquire antigen presentation capabilities and can activate antigen-specific T cells *in vitro*. These results provide detailed analyses of the membrane-coated nanoparticle generated from extrusion, sonication, and a new combined process, and highlight the antigen-specific therapeutic potential of DCmPs, and have broad implications for the development of other membrane-coated particle systems.

## 2. Materials and methods

### 2.1. Cell Culture

DC2.4 cell line (Merck, Cat. # SCC142) were cultured in RPMI 1640 medium (Gibco, Cat# 21870076) supplemented with 10% Fetal Bovine Serum (FBS) (Gibco, Cat# A5670801), 1x Penicillin-Streptomycin (P/S) (Gibco, Cat# A5873601), 1x L-glutamine (Gibco, Cat# 25030081), 1x MEM NEAA (Gibco, Cat# 11140050), 1x (or equivalent to 25 mM) HEPES (Corning, Cat# 25060CI), and 0.0054x β-Mercaptoethanol (Gibco, Cat# 21985023). B3Z was kindly gifted by Dr. James Moon at University of Michigan. Cells were cultured in RPMI 1640 medium supplemented with 10% FBS, 1x P/S, 1mM sodium pyruvate (Gibco, Cat# 11360070), and 55µM β-Mercaptoethanol. DOBW cells were generously gifted by Dr. Clifford Harding at Case Western Reserve University. DOBW cells were cultured in RPMI 1640 medium supplemented with 10% FBS, 1x P/S, 1mM sodium pyruvate, 1x HEPES, and 15µM β-Mercaptoethanol at 37°C. All cells were maintained in a humidified incubator with 5% CO₂.

### 2.2. BMDC Differentiation

Bone marrow (BM) cells were harvested from femurs and tibias of C57BL/6 mice obtained following Syracuse University IACUC protocol (P3-24). After RBC lysis, BM cells were washed and passed through a 70 μm cell strainer to obtain a single-cell suspension. The cells were then resuspended at a density of 2 × 10⁶ cells/mL in 6-well plate and cultured in RPMI 1640 medium supplemented with 10% FBS, 2 mM L-glutamine, 10 mM HEPES, 50 μM β-mercaptoethanol, and 1× P/S, in the presence of 200 ng/mL recombinant murine Flt3-ligand (PeproTech, Cat# 25031L100UG). On day 3, 60% of the culture medium was gently replaced with fresh Flt3L-containing medium. Immature BMDCs were harvested by gentle pipetting for downstream experiments on day 7.

### 2.3. Adjuvant Stimulation and Cytotoxicity

DC2.4 cells were stimulated with ovalbumin (OVA) protein (Sigma, Cat# A55031G) at a concentration of 0.3 mg/mL in combination with different adjuvants, including lipopolysaccharide (LPS) (Norus Biologicals, Cat# NBP2252951mg) at 50, 100, 200, or 400 ng/mL, Poly(I:C) (Fisher Scientific, Cat# NC9180242) at 25, 50 100, or 200 μg/mL, and interferon-gamma (IFN-γ) (Fisher Scientific, Cat# 485MI100CF) at 1.25, 2.5, 5, or 10 μg/mL. The stimulation was carried out overnight in 96-well plates, with each well containing 2 × 10^5^ cells in 200 µL of the respective treatment conditions. The cytotoxicity of adjuvant treatment was evaluated using the CellTiter 96® AQueous One Solution Cell Proliferation (MTS) Assay (Promega, Cat# G3582), following the manufacturer’s instructions. Briefly, 20 μL of the assay reagent was added to each well and incubated at 37 °C for 1-4 hours, after which the absorbance was measured at 490 nm using a 96-well plate reader (Synergy/H1, BioTek). Cell viability was calculated and expressed as a percentage using the formula: 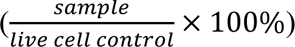.

### 2.4. DC2.4 Membrane Isolation

The DC2.4 cells were lifted using cell scrapers and washed and resuspended in 1× Tris buffer (Thermo Scientific, Cat# J60764K2). Subsequently, the cells were resuspended in 1× Tris buffer supplemented with EDTA and a protease inhibitor cocktail (Thermo Scientific, Cat# 78442) for membrane isolation (Figure S5). The cells were mechanically disrupted using a Dounce homogenizer with a loose-fitting pestle for 40 times, and a tight-fitting pestle for 40 additional passes. The homogenized lysate was centrifuged at 3000 × g for 10 minutes, followed by a second round of homogenization. The supernatant after both centrifugations were combined in a centrifuge tube (Beckman Coulter, Cat# 344059) and ultracentrifuged at 100, 000 × g for 60 minutes at 4 °C. The pellet, cell membrane proteins, was resuspended in 1 × PBS containing 0.05% TWEEN-20. Finally, the protein concentration was measured using either the Bradford (Thermo Scientific, Cat# A55866) or BCA (Thermo Scientific, Cat# 23225) assay and stored at -80°C. We approximately recover 1.5 mg of membrane protein per 100 million DC2.4 cells. For CFSE-labeled membrane proteins, we first label DC2.4 cells with CellTrace CFSE staining solution (Invitrogen, Cat# C34554) following the manufacturer’s instructions before harvesting cells in hypotonic lysing buffer for homogenization.

### 2.5. Fabrication And Characterization of DC Membrane-Coated Nanoparticles (DCmPs)

The poly(lactic-co-glycolic acid) (PLGA) nanoparticles (NPs) were prepared via nanoprecipitation by adding 1 mg/mL PLGA (Polysciences, Cat# 23986-5) acetonitrile solution into 10 mL of 1% polyvinyl alcohol (PVA) (Sigma, Cat# 363170-25G) dropwise under rapid stirring (600 rpm) under room temperature. The stirring was reduced to 200 rpm after 4 hours to let organic solvent evaporate overnight. If fluorescent labeling was required, rhodamine B (RhoB) (Thermo Scientific, Cat# A13572.18) was added to 1% PVA solution at a final concentration of 0.005% w/v before initiating the precipitation process. The following day, the nanoparticles were washed five times using deionized water and a Centricon 100 kDa filter (Millipore, Cat# UFC910024) at 4000 × g for 10 minutes. Then, the nanoparticles were collected after lyophilization for coating.

To fabricate DCmPs, various coating processes were used, including sonication, extrusion and a combined sonication-extrusion process (Figure 2A). In the sonication process, a mixture of nanoparticles and membrane proteins were prepared at a 1:1 weight ratio and sonicated for 5 minutes. Unbound proteins were removed via centrifugation at 10,000 × g for 10 minutes. In the extrusion process, nanoparticles and membrane proteins were separately prepared in an extruder set (Avanti, Cat# 610000-1EA) at 1:1 ratio. Then, nanoparticles and membrane proteins were mixed by 11 extrusion passes through a 200 nm filter (Cytiva, Cat# 10417004) and drain discs (Cytiva, Cat# 230300), followed by the removal of unbound proteins using centrifugation at 10,000 × g for 10 minutes. In the combined sonication-extrusion process, the mixture of nanoparticles and membrane proteins at 1:1 ratio was prepared and sonicated for 5 minutes. After the unbound proteins were removed using centrifugation at 10,000 × g for 10 minutes, the obtained particles were resuspended and transferred into extruder for mixing 11 passes through extrusion. The efficiency of membrane coating on PLGA nanoparticles was evaluated using the Bradford assay. These protein quantification methods were employed to determine the amount of membrane proteins successfully adsorbed onto the nanoparticle surface. Quantification was calculated and expressed as a percentage using the formula: 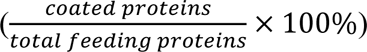.

### 2.6. Flow Cytometry Analysis of DC2.4 Surface Markers

Cells were firstly stained with LIVE/DEAD Fixable Dye, namely FVD (Invitrogen, Cat# L34962A) for 20 minutes at room temperature (RT), followed by staining surface proteins using fluorophore-conjugated antibodies specific for MHC class II (MHC-II, APC-eFluor 780) (Invitrogen, Cat# 47532182), MHC class I (MHC-I, PerCP-eFluor 710) (Invitrogen, Cat# 46599882), SIINFEKL/H-2Kb (25-D1.16, BV711, BD, Cat# 756314), CD80 (Brillian Violet 605) (BD, Cat# 563052), CD86 (PE-Cy7) (BD, Cat# 560582), and ICAM-I (BV711) (BD, Cat# 753781) in the presence of Fc blocking 2.4G2 antibody (BD, Cat# 553142) for 20 minutes at RT. The stained cells were washed twice with 1% FBS-containing flow buffer and fixed with 50 µL of 4% paraformaldehyde solution (Thermo Scientific, Cat# J61899AP) for 10 minutes at RT. For Eα/MHC-II Presentation by DC2.4 cells or BMDCs, cells were treated with Eα_(52-68)_ peptide (ASFEAQGALANIAVDKA, GenScript) in the presence of LPS before stained similarly as mentioned above using Ea52-68 peptide bound to I-Ab Antibody (eBioY-Ae, FITC, eBioscience). DCmP particles were similarly stained using antibodies but was centrifuged at 10,000 × g for 10 minutes during washing. The final samples were resuspended in 200 µL of flow buffer for flow cytometry analysis using the BD LSR Fortessa flow cytometer, and results were analyzed using FlowJo software (version 10.10).

### 2.7. Dynamic Light Scattering (DLS) Measurement

DLS measurements were performed using a Zetasizer instrument operated via the ZS Xplorer software. Samples were prepared at a concentration of 1 mg/mL or diluted accordingly and measured using the DTS0012 or DTS1070 cuvette for determining the average particle size distribution and zeta potential. The polystyrene latex as the dispersant was configured for measurement parameters. A single read per sample, a measurement temperature of 25 °C, and an equilibration time of 120 seconds were set.

### 2.8. Transmission Electron Microscopy (TEM) Imaging

The samples were prepared at ≥ 10 mg/mL concentration in 1 × PBS. Four microliters of the sample was drop-cast onto a carbon coated TEM grid for 30s @ 15mA to promote particle adsorption. Sample was adsorbed for 5 minutes, then excess liquid wicked with filter paper. Grids were washed with UltraPure distilled water 2 × for 1-2 minutes each. Then, 1% Uranyl acetate was stained for 1 minute, excess wicked away, grids allowed to dry on filter paper for a few minutes and then placed in a grid box and held in a vacuum desiccator until imaging. Imaging was conducted with a JEOL JEM-1400 TEM operated at 80kV, and digital micrographs were acquired with a Gatan Orius SC1000 CCD camera. The obtained TEM images were further processed and analyzed using ImageJ software.

### 2.9. SDS-PAGE and Western Blotting

Protein samples were prepared by mixing the desired amount of protein with 2 × Sample Buffer (Novex, Cat# LC2676) in a 1:1 ratio (v/v). The samples were then heated at 75°C for 5 minutes and loaded into the wells of 4-12% Tris-Glycine Gel (Invitrogen, Cat# XP04120BOX) using 1 × Running Buffer (Invitrogen, Cat# NP0001). Five microliters of the protein ladder (Thermo Scientific, Cat# 26619) were also loaded alongside the protein samples before running the gel. After completion, the gel for SDS-PAGE analysis was stained with Blue Stain (Thermo Scientific, Cat# 24590) for 1 hour on a shaker.

For Western blotting analysis, gel was transferred onto 0.45μm nitrocellulose membrane at 75 V for 75 minutes. The membrane was then blocked with 5% skim milk in TBST (TBS containing 0.05% TWEEN20) for 1 hour at RT. The membrane was then incubated with the primary antibodies (MHC class II (MHC-II) (Invitrogen, Cat# PA5116820), MHC class I (MHC-I) (Invitrogen, Cat# MA548095), CD80 (Invitrogen, Cat# PA579001), CD86 (Invitrogen, Cat# MA535211), ICAM-I (Invitrogen, Cat# 701254), and Alpha Na^+^/K^+^ ATPase (Invitrogen, Cat# ANP001-200UL) in 2% skim milk in TBST for either 2 hours at RT or overnight at 4°C. The membrane was then washed 4-5 times with TBST before incubating it with the secondary antibody (HRP-conjugated Ab) (Thermo Scientific, Cat# 31460) in 5% skim milk in TBST for 1 hour at RT. After another 4-5 washes with TBST, proceed with protein detection using a ChemiDoc^TM^ MP Imaging System (Bio-Rad). Quantitative analysis of Western blot protein bands using Fiji (ImageJ) software. Band intensities were normalized to Na⁺/K⁺-ATPase, and fold changes were calculated relative to appropriate controls.

### 2.10. Interaction Between Immune Cells and DCmPs

For flow cytometry, 2 × 10⁵ DC2.4, B3Z, or DOBW cells were seeded into 96-well plates and treated with each NP formulation at a final concentration of 100µg/mL (based on PLGA weight) for 4 hours at 37°C with 5% CO₂ in five biological replicates. Cells were washed three times with cold flow buffer, then stained with FVD for 10 minutes at RT before fixing with 4% paraformaldehyde. Flow cytometric analysis was performed using a BD LSRFortessa™ analyzer.

For confocal microscopy, 2 × 10⁵ DC2.4 cells in sterile 4-well chamber slides (SPL, Cat# 30104) were treated with the respective NPs at a final concentration of 100µg/mL (based on PLGA weight) for 4 hours at 37°C with 5% CO₂. The cells were gently washed with PBS containing 0.05% Tween-20 (PBST) twice to remove unbound NPs, then fixed with 4% paraformaldehyde for 10 minutes at RT and subsequently stained with DAPI (Thermo Scientific, Cat# 202710100) to visualize nuclei. Coverslips were mounted onto glass slides using Permount Mounting Media (Fisher Chemical, Cat# SP15-100). Imaging was performed using a Leica Dmi8 confocal laser scanning microscope (CLSM). The co-localization analysis of RhoB-labeled PLGA and CFSE-labeled membrane was conducted using Fiji (ImageJ) following Manders’ coefficients: M1 for proportion of RhoB-labeled PLGA signal overlapping with CFSE-labeled membrane while M2 for proportion of CFSE-labeled membrane signal overlapping with RhoB-labeled PLGA. The resulting Manders’ coefficients were multiplied by 100 to express co-localization as a percentage. Four independent images were used for analysis (Figure S11).

### 2.11. DOBW Cell Activation Assay

DOBW cells were seeded at 2 × 10⁵ cells per well in a 96-well plate and cocultured with 100 μg/mL DCmP or nonstim DCmP, or with OVA/LPS-stimulated DC2.4 cells or BMDCs at 1:1 ratio. As a negative control, 2 × 10⁵ DOBW cells were cultured alone or with no stimulated DC2.4 or BMDCs cells. All cultures were incubated at 37°C with 5% CO₂ for 3 days, before the IL-2 concentrations within media were assessed using an ELISA kit (Invitrogen, Cat# BMS601), according to the manufacturer’s instructions.

### 2.12. B3Z Cell Activation Assay

B3Z cells were seeded at 2 × 10⁵ cells per well in a 96-well plate, followed by the addition of DCmPs or nonstim DCmPs at a concentration of 100 μg/mL. For comparison, 2 × 10⁵ immature or OVA/LPS-stimulated DC2.4 cells were added to B3Z cells for 2-day coculture. After the 2-day incubation, cell media were replaced with 150 μL of CPRG/lysis buffer, composed of 0.15 mM chlorophenol red-β-D-galactopyranoside (CPRG) (Sigma, Cat# 59767-25G-F), 0.1% Triton X-100, 9 mM MgCl₂, and 100 μM β-mercaptoethanol in PBS. The cells were then resuspended and incubated at 37 °C in the dark for 90 minutes. The absorbance at 570 nm was determined using a microplate reader (Synergy/H1, BioTek).

### 2.13. Statistical Analysis

Data were analyzed using GraphPad Prism version 10.4.2. Statistical comparisons between two groups were performed using an unpaired Student’s t-test, while comparisons among three or more groups were conducted using one-way ANOVA followed by Tukey’s multiple comparison test.

## 3. Results

### 3.1. Characterization of the Cell Membrane of Stimulated DC2.4 Cells

DCs orchestrate the initiation of adaptive immune responses and activate T cells through antigen presentation.^24^ This antigen presentation process relies critically on many DC membrane proteins, including pMHC to engage cognate T-cell receptor (TCR), CD80/CD86 as costimulatory molecules, and ICAM-1 as adhesive molecules to facilitate the formation of DC-T cell immune synapses.^25^ DCmPs can leverage these membrane proteins to mimic DC function to engage antigen-specific T cells and mediate antigen presentation.^8,21–23^ To maximize the surface protein expression by DC2.4 cells and facilitate the subsequent protein coating processes, we compared several common stimulation conditions for the surface protein upregulation, including LPS (agonist for Toll-like receptor 4, TLR-4), Poly(I:C) (TLR-3 agonist), and IFNγ (DC-activating cytokine).^35,36^ (Figure S1, S2) After 12 hour stimulation in the presence of OVA protein antigens, we determined the DC2.4 surface protein level using flow cytometry. We found that LPS promoted high level of MHC class I (MHCI) and CD80 within the tested concentration range, while IFNγ stimulation leads to the highest expression of ICAM-1 and CD86. (Figure S1) All stimulated conditions showed minimal cytotoxicity. (Figure S2) Considering the relatively low amount of LPS (50ng/mL) required to achieve high level of membrane protein upregulation and the considerably higher cost associated with other adjuvant options, we elect to focus on determining protein expression in DC2.4 cells stimulated with LPS and OVA in subsequent studies. (Figure 1A) Notably, although LPS unregulated the expression of MHC Class II (MHC-II) in DC2.4, the overall MHC-II proteins expression level was at a very low level, consistent with other study.^37^ (Figure S3) We further determined the overall protein expression profile and the level of presentation-related membrane proteins in the lysates of LPS-stimulated and non-stimulated DC2.4 cells using SDS-PAGE and western blot. SDS-PAGE revealed a more complex and intensified protein banding pattern in stimulated DC2.4 cells relative to non-stimulated cells, indicating enhanced protein expression following LPS treatment. (Figure 1B) Western blot analysis further confirmed the elevated expression of all examined key presentation-related membrane proteins in LPS-stimulated DC2.4, consistent with flow cytometry experiments. (Figure 1C) The expression fold-changes were quantified by normalizing against membrane protein control (Na^+^/K^+^ ATPase), and showed that LPS-stimulation improved DC2.4 expression of MHCI by 4.5 fold, CD80 by 1.58 fold, CD86 by 2.25 fold, and ICAM-1 by 1.16 fold relative to non-stimulated cells. (Figure S4) These results confirm that LPS stimulation effectively promotes presentation-related membrane protein expression in DC2.4 cells, thereby potentially benefiting downstream membrane protein isolation and nanoparticle coating. Therefore, in subsequent experiments, DC2.4 cells were all stimulated with both OVA antigens and LPS adjuvant, unless specified otherwise.

**Figure 1:**
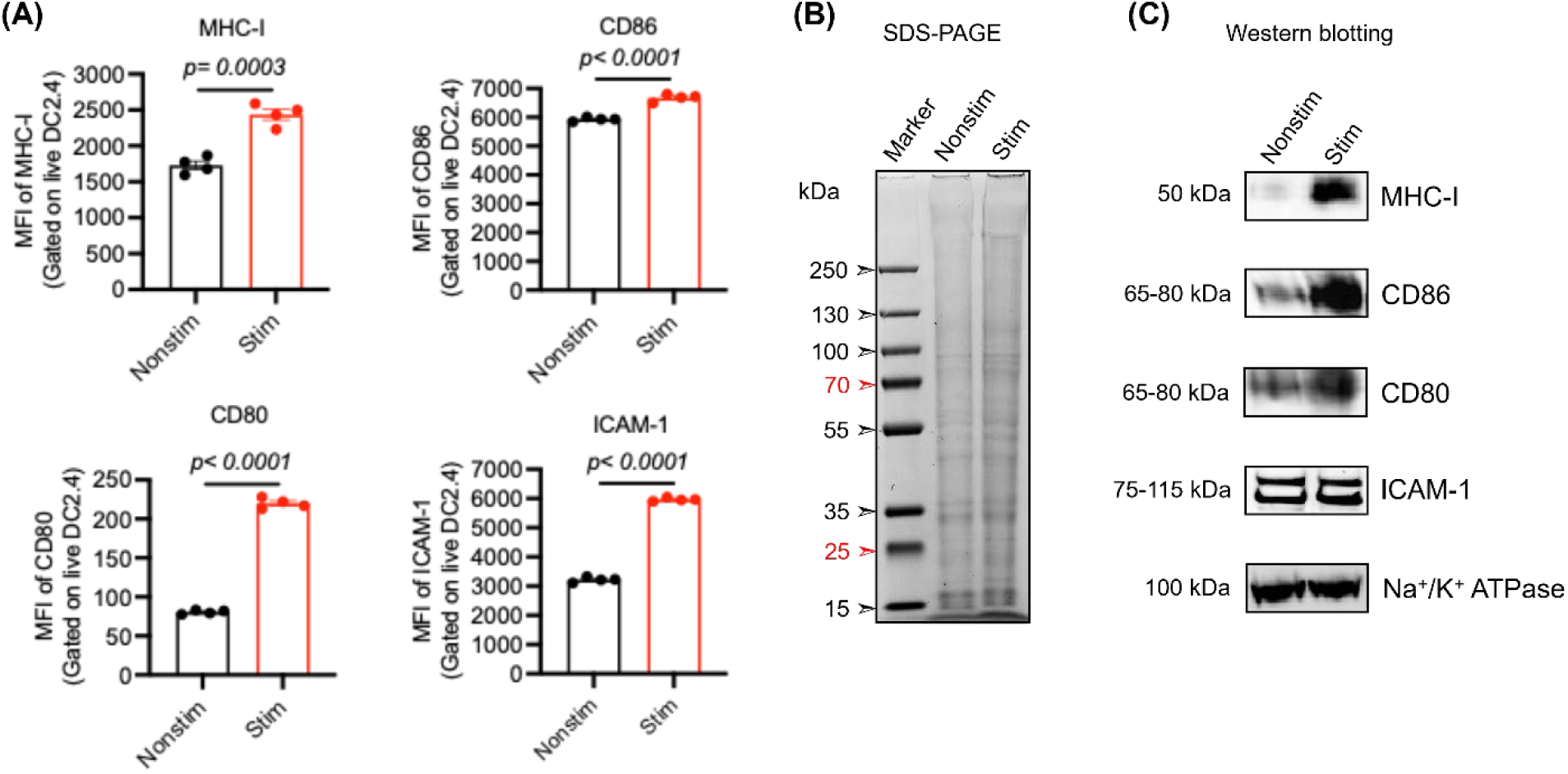
LPS stimulation enhances the expression of antigen presentation–related membrane proteins in DC2.4 cells. DC2.4 cells were stimulated with OVA protein (0.3 mg/mL) and LPS (50 ng/mL) for 15 hours. The activation status of the cells was assessed by evaluating the expression of MHC class I (MHC-I), CD80, CD86, and ICAM-1 via flow cytometry and western blotting. **(A)** Flow cytometry analysis showing the surface expression of activation markers in non-stimulated (Nonstim) versus stimulated (Stim) DC2.4 cells.(N=4) **(B, C)** SDS-PAGE and western blot analyses comparing protein expression between Nonstim and Stim groups. Statistical significance was determined using Student’s *t*-test.

### 3.2. Fabrication and Physical Characterization of DCmPs

To produce membrane-coated particles, we first isolated the membrane protein of LPS-stimulated DC2.4 through homogenization and ultracentrifugation. The protein extraction was confirmed using SDS-PAGE gels at each step. (Figure S5) PLGA nanoparticles with an average diameter of 100 nm were manufactured through nanoprecipitation as the core for membrane coating.^38^ (Figure 2B) To produce DCmPs, we adopted two conventional coating approaches: sonication and extrusion. Furthermore, we developed a combined approach, extrusion following sonication, for membrane coating. (Figure 2A) We hypothesize that the additional extrusion process following sonication will improve the diameter uniformity of the obtained DCmP without reducing coated protein amount. We first compared the diameters and zeta potentials of the obtained coated nanoparticles from two separate experiments with 5 individually prepared samples for each coating approaches per experiment. The average diameters of the coated particles for each approach respectively are 340 nm for sonication, 350 nm for extrusion, and 180 nm for combined group, all larger than the uncoated bare PLGA NPs (100 nm). (Figure 2B-C) The increase in particle hydrodynamic diameters suggests the presence of protein coating in the obtained particles. The polydispersity index (PDI) of DCmPs from the three coating processes were also determined using DLS and both sonication- and extrusion-produced DCmPs showed broader size distribution relative to bare NPs (Figure 2D) The additional extrusion process in combined coating process reduced size distribution, indicating improved uniformity and reproducibility. It is worthnoting that the centrifugation process (10,000 × g, 10 mins) for unbound protein removal resulted in aggregation of the final products, for the diameter and PDI of particles are lower for both sonication and extrusion processes before centrifugation.(Figure S6) The negative zeta potential of all the DCmP groups relative to bare NPs also indicate the presence of protein coating, as confirmed in other studies.^3,4^ (Figure 2E) The presence of protein coating was confirmed using Transmission electron microscopy (TEM). A distinct membrane layer can be found on the surface of DCmPs fabricated by all three approaches. (Figure 2F) These results confirm the successful incorporation of stimulated DC2.4 membrane onto PLGA NPs. Among the three coating strategies, the combination of sonication and extrusion approach yielded the DCmP with the lowest diameter and size distribution. Protein coatings can be found on DCmP from all three coating approaches.

**Figure 2.**
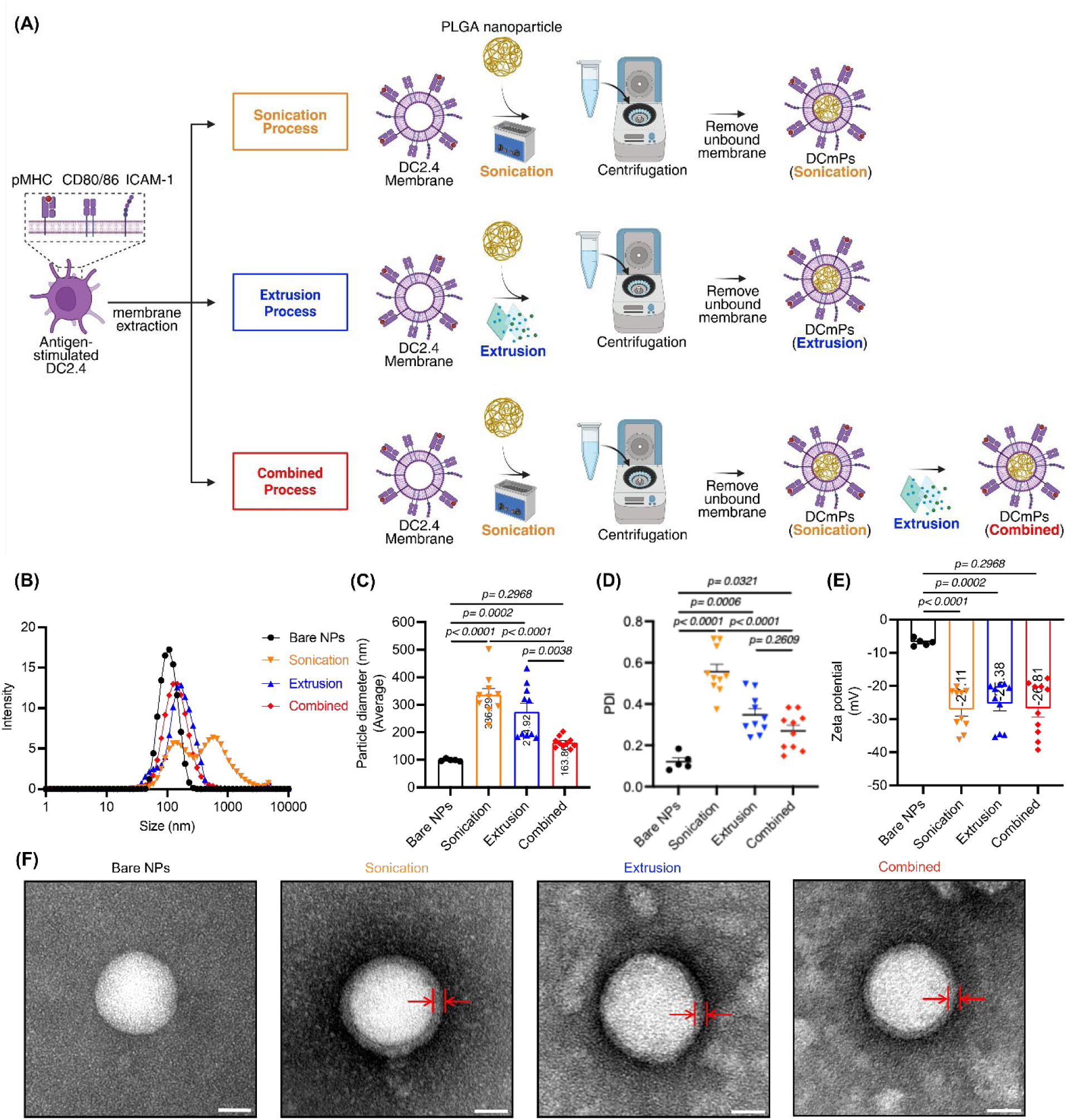
DCmPs fabricated via the combined approach show the lowest diameter and the most uniform size distribution relative to sonication and extrusion alone. **(A)** Schematic illustration of the fabrication process for DCmPs. Cell membranes of DC2.4 stimulated with OVA protein (0.3 mg/mL) plus LPS (50 ng/mL) were isolated using Dounce homogenization followed by ultra-centrifugation and subsequently coated onto PLGA NPs using one of three methods: sonication, extrusion, or a combined process (sonication followed by extrusion) at a 1:1 weight ratio (w/w). Unbound proteins were removed by centrifugation, and the resulting purified DCmPs were characterized using dynamic light scattering (DLS) and transmission electron microscopy (TEM). The final products are analyzed via DLS for **(B)** size distribution, **(C)** average particle size, **(D)** polydispersity index (PDI), and **(E)** zeta potential of bare nanoparticles (Bare NPs) and DCmPs fabricated using different coating processes. N=10 **(F)** TEM images confirming the membrane coating of DCmPs prepared by the respective processes. Scale bars represent 20 nm. Statistical significance was determined using one-way ANOVA followed by Tukey’s multiple comparison test.

### 3.3. Protein Characterization of DCmPs

Having confirmed the presence of protein coating, we proceed to determine the amount of proteins on coated particles using Bradford assay and calculate protein coating efficiency 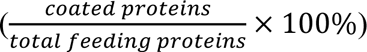. Using 500µg/mL feeding protein concentration at 1:1 ratio (PLGA NP : membrane protein), DCmPs produced via sonication alone were coated with around 15 % of total feeding proteins, and DCmPs from combined approach showed around 11% protein coating efficiency. (Figure 3A) Extrusion alone, however, showed significantly lower protein coating efficiency relative to sonication, and only achieved 5% coating efficiency. The low coating efficiency by extrusion was further confirmed by the fact that significantly higher amount of unbound proteins were found in the supernatant of extrusion samples than in the sonication samples. (Figure S7) We subsequently assessed the protein composition of the DCmPs via SDS-PAGE and western blot gels. According to SDS-PAGE gel, DCmPs from the three coating processes all showed the protein profile closely resembling the isolated DC2.4 cell membrane protein, suggesting that all three approaches can preserve the overall membrane protein composition of the membranes. (Figure 3B) The SDS-PAGE analysis of the supernatant (unbound proteins removed during centrifugation) showed distinct protein profile, where protein bands were largely absent with the exception of protein with a molecular weight between 70 kDa and 130 kDa, suggesting successful and largely unbiased membrane protein coating on DCmPs. (Figure 3B) Using western blot, we also successfully identified the presence of key presentation-related membrane proteins on DCmPs, including MHC-I, co-stimulatory molecules (CD86 and CD80), and the adhesion molecule (ICAM-1) in DCmPs from all three coating approaches (Figure 3C). The band intensity of these proteins was quantified based on Na^+^/K^+^ ATPase control, but the impact of coating approaches on the retention of these individual proteins is unclear. (Figure S8) Taken together, all three coating approaches similarly maintain the overall membrane protein composition on the protein coating and preserve the key DC membrane proteins on DCmPs. Furthermore, the sonication and combined approach yielded superior protein coating efficiency, relative to extrusion alone.

**Figure 3.**
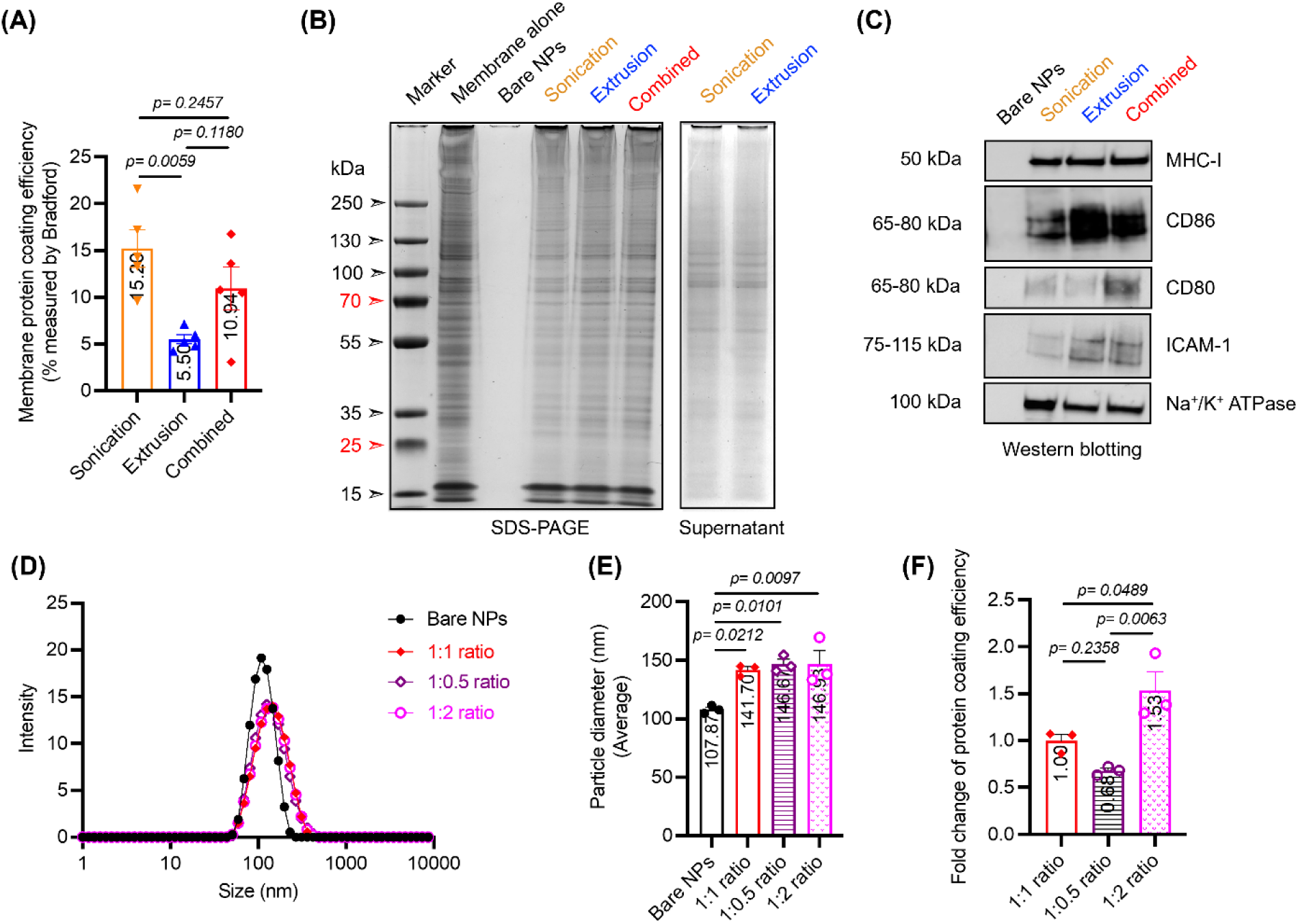
DCmPs retain key DC membrane proteins. DCmPs were fabricated using sonication, extrusion, or the combined approaches. **(A)** The membrane coating efficiency of different approaches were assessed and compared using Bradford assay.(N=5) **(B)** SDS-PAGE analysis confirm the similarity of the protein profiles between isolated DC membrane proteins and DCmPs (left panel) and show the unbound proteins in the supernatant after centrifugation (right panel). **(C)** Western blot analysis confirm the presence of key membrane proteins on DCmPs fabricated via different processes at a 1:1 ratio (w/w). **(D–F)** Characterization of DCmPs produced using the combined process at different nanoparticle-to-protein ratios (1:1, 1:0.5, or 1:2, w/w), N=3 **(D)** size distribution by DLS, **(E)** average particle size by DLS, and **(F)** fold-change in membrane coating efficiency. Statistical significance was determined using one-way ANOVA followed by Tukey’s multiple comparison test.

As higher coated protein amount can potentially benefit downstream application, we examined the protein coating efficiency of the combined approach at three different weight ratios, 1:0.5, 1:1, 1:2 (PLGA NP : membrane protein) to potentially improve the amount of coated proteins. Increasing feeding protein amount indeed increased the amount of coated proteins, without significant alteration in particle diameters likely thanks owing to the additional extrusion process of the combined approach. (Figure 3D, 3E, 3F) But we did not achieve 2-fold increase in the coating efficiency when feeding protein amount was doubled, indicating 1:1 ratio is potentially close to the saturation point for protein coating based on current design. (Figure 3F) Given that a 1:1 weight ratio between PLGA NP and membrane protein has demonstrated consistency in both the size and protein coating of the resultant DCmPs, we decided to use this ratio for DCmPs production for all the following experiments in order to minimize proteins usage.

### 3.4. Characterization of the Composition of DCmPs

Having confirmed the presence of membrane proteins, including the key membrane proteins involved in DC antigen presentation, we sought to gain a more detailed insight regarding the composition of DCmPs produced by the three approaches. It is well established that the homotypic interaction promotes binding between membrane-coated nanoparticles and membrane source cells.^2,3,10^ Thus, DCmPs and empty DC2.4 cell membranes should both be readily taken up by DC2.4 cells due to this homotypic interaction. Because the compositions of nano-scale particle samples are difficult to study directly using flow cytometry or confocal microscopy, we propose to leverage this interaction and use DC2.4 cells as a surrogate to detect the presence of DCmPs and empty membrane vesicles. To this end, we produced Rhodamine B (RhoB) -encapsulated PLGA NPs and coated the RhoB+ cores with DC2.4 membrane proteins labeled with carboxyfluorescein succinimidyl ester (CFSE) via the three different coating approaches. The diameters and the zeta potentials of the obtained RhoB/CFSE-labeled DCmPs were found to be consistent with previous results. (Figure S9) We hypothesize that DC2.4 cells that have internalized DCmPs will be positive for both RhoB and CFSE, and DC2.4 cells that take up only PLGA NPs or membrane proteins will be positive for RhoB or CFSE respectively. After a 4-hour incubation between DC2.4 cells and DCmPs, flow cytometry revealed that approximately 80% of cells have internalized both CFSE and RhoB, while relatively low percentage of DC2.4 are positive for single fluorophore with 4-9% for CFSE and 12-14% for RhoB respectively. (Figure 4A, 4B, 4C, 4D) DC2.4 internalization of DCmPs was consistent amongst the three coating strategies. This flow cytometry result suggests that about 80% of particles are protein-coated particles, roughly 14 % PLGA NPs are uncoated, and low percentage of proteins present are unbound proteins. The combined approach appears to yield lowest amount of unbound protein for only 4.7% DC2.4 are CFSE+, relative to 9.4% in sonication group and 8.5% in extrusion. (Figure 4B) To confirm the double positive DC2.4 did not just take up a mixture of PLGA NPs and membrane proteins, we treated DC2.4 cells with a mixture of CFSE-labeled DC2.4 membrane proteins and RhoB-PLGA NPs, in addition to protein alone or NP alone control. (Figure S10) The membrane protein alone group showed robust uptake by DC2.4 cells as expected, with 84% of DC2.4 cells positive for CFSE. (Figure S10B, S10D) Incubation with the membrane and NP mixture only yield about 1% of DC2.4 that are positive for both CFSE and RhoB, far lower than the 80% frequency observed in DCmP treatment.(Figure S10C, S10D) Therefore, we conclude that the double-positive population observed in DCmP treated DC2.4 cells is likely the result of DCmP-binding, but not caused by the internalization a mixture of membrane proteins and NPs.

**Figure 4.**
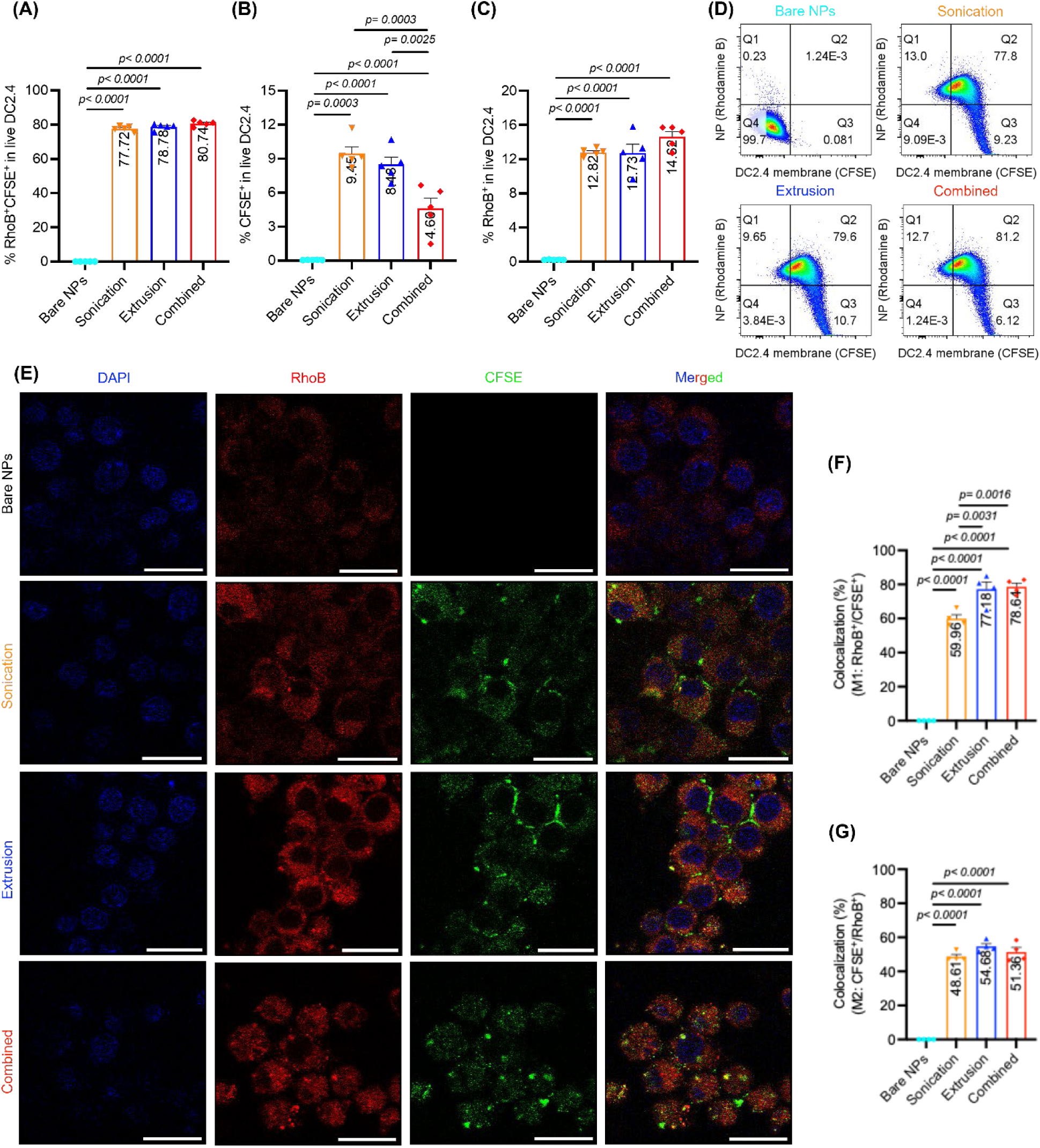
The characterization of DCmP compositions using DC2.4 cells as the surrogate. DCmPs were formed using CFSE-labeled DC2.4 membrane proteins and RhoB-labeled PLGA NPs and were incubated with DC2.4 cells for 4 hours before the compositions were determined using flow cytometry and confocal laser scanning microscopy (CLSM). **(A–D)** Flow cytometry analysis show the frequencies of DC2.4 cells positive for **(A)** RhoB (PLGA NPs) signal, **(B)** CFSE (membrane protein) signal, **(C)** dual positivity (both PLGA NPs and membrane proteins), and **(D)** representative flow cytometry plots from five independent experiments. (N=5) **(E)** Representative CLSM images of the cellular uptake and intracellular localization of DCmPs in DC2.4 cells from four independent experiments (Figure S11). The co-localization frequencies were quantified using Manders’ coefficients via Fiji (ImageJ) software: **(F)** M1 for proportion of RhoB-labeled PLGA NPs overlapping with CFSE-labeled membrane; **(G**) M2 for proportion of CFSE-labeled membrane signal overlapping with RhoB-labeled PLGA NPs. N=4, Scale bars: 20 μm. Statistical significance was determined using one-way ANOVA followed by Tukey’s multiple comparison test.

To analyze the protein coverage of DCmPs with greater detail, we imaged DC2.4 cells after 4h treatment with fluorophore-labeled DCmPs using CLSM. (Figure 4E, 4F, 4G, Figure S11) Through flow cytometry, we found nearly all cells in the field of view are positive for both RhoB and CFSE, which is consistent with the observed high frequency (about 80%) of (RhoB+CFSE+) DC2.4 cells. We assume that the colocalization between RhoB (Red, PLGA NPs) and CFSE (green, membrane proteins) indicates nanoparticles coated with proteins. We chose the Manders’ coefficients as the statistical parameters to determine the proportion of CFSE signal overlapping with RhoB (the frequency of PLGA NPs that are coated with membrane proteins) and the proportion of RhoB signal overlapping with CFSE (the frequency of proteins associated with NPs). Approximately 60-79 % NPs are colocalized with proteins, indicating that the majority of PLGA NPs are coated with proteins and about 20-40% of NPs are uncoated. (Figure 4F) Notably, both extrusion and combined approaches yielded slightly higher protein coverage on PLGA NPs relative to sonication alone, indicating extrusion process is potentially more effective in promoting proteins and NP interactions. Additionally, about 49-55% of proteins were found to colocalize with PLGA NPs, suggesting that nearly half of protein present are unbound despite of our best effort to remove unbound proteins. (Figure 4G) Confocal imaging largely corroborates with the flow cytometry experiments, indicating that majority of PLGA NPs are coated with membrane proteins in the three groups and that unbound proteins are present in all the samples.

Significant discrepancies exist in the composition of DCmPs when determined using flow cytometry and confocal imaging. These differences may be attributed to several factors, including the sampling bias in confocal imaging and its ability to provide a more detailed view of intracellular particles compared to flow cytometry. Taken together, DCmPs produced via the three coating strategies are all similarly comprised of a significant number of membrane-coated particles, along with relatively low percentage of bare NPs and unbound proteins.

### 3.5. Antigen-Specific T Cell Binding by DCmPs

Having confirmed the presence of key presentation-related proteins, including MHC-I, CD80/86, ICAM-1 on DCmPs, we sought to determine whether these membrane proteins can facilitate the binding between DCmPs and T cells. We first confirmed the SIINFEKL/MHC-I presentation on OVA/LPS-stimulated DC2.4 cells using 25-D1.16 antibodies via flow cytometry. (Figure 5A) Given that DC2.4 cells do not express significant levels of MHC-II proteins, (Figure S3) we further determined the MHC-II antigen presentation capability in DC2.4 cells by treating DC2.4 with Eα_52–68_ peptides (50 µM) in the presence of LPS for 12 hours. Indeed, we found no detectable level of Eα_52–68_/MHC-II presentation in DC2.4 cells using Y-Ae antibodies, unlike bone marrow-derived dendritic cells (BMDCs) that showed Eα_52–68_/MHC-II presentation after the same treatment. (Figure S12) Therefore, we conclude that DC2.4 cells exclusively present MHC-I antigens, as demonstrated by others.^37^

**Figure 5.**
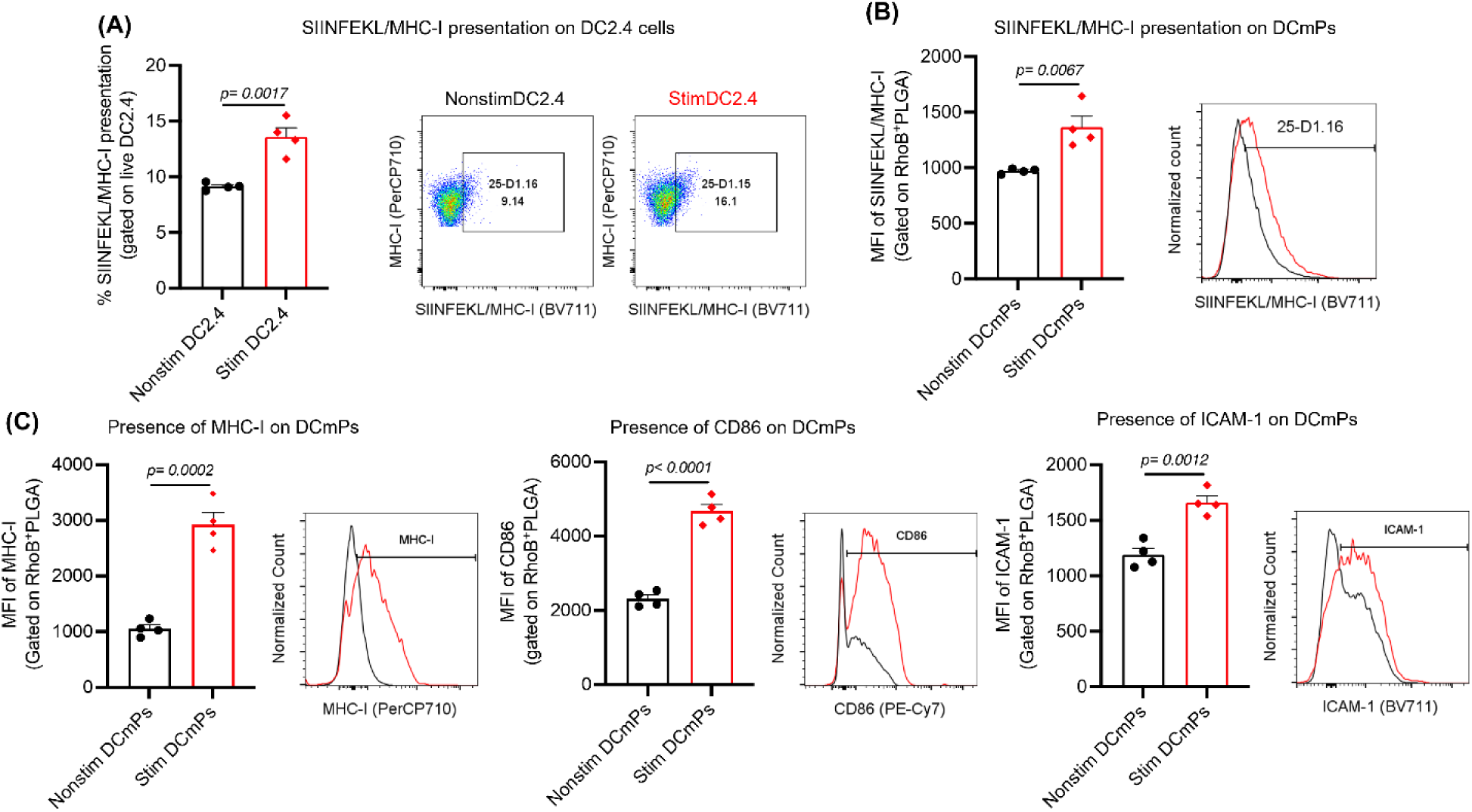
Stimulation of DC2.4 cells improves presentation–related membrane proteins carried by DCmPs. **(A)** Flow cytometry confirmed the upregulation of SIINFEKL/MHC-I complex in DC2.4 cells after the stimulation with OVA protein (0.3 mg/mL) and LPS (50 ng/mL). **(B–C)** Comparison of antigen presentation–related membrane proteins displayed on DCmPs. Membranes from non-stimulated or stimulated DC2.4 cells were used to coat Rhodamine B (RhoB)-labeled PLGA NPs via the combined approach. Flow cytometry analysis was performed on the RhoB⁺ gated population to assess surface protein expression. Stimulation of DC2.4 cells improved the amount of **(B)** SIINFEKL/MHC-I complex, **(C)** MHC-I molecules, CD86, and ICAM-1 on DCmPs. N=4, Statistical significance was determined using Student’s *t*-test.

Because non-stimulated DC2.4 cells have a lower level of protein expression relative to LPS-stimulated cells, we sought to confirm that PLGA NPs coated with the membrane proteins from non-stimulated DC2.4 cells will have lower level of presentation-related proteins. We prepared these particles using the combined coating method, and they will be referred to as nonstim DCmPs for brevity hereafter. As expected, DCmPs displayed significantly higher levels of the three examined surface proteins, i.e. MHC-I, CD86, and ICAM-1, according to flow cytometry analysis. We also confirmed the presence of higher level of SIINFEKL/MHC-I on DCmP as compared to nonstim DCmPs. (Figure 5B, 5C) We hypothesize that the elevated protein levels on DCmPs can promote particle binding to antigen-specific T cells, relative to nonstim DCmPs. Moreover, since DC2.4 cells exclusively present SIINFEKL/MHC-I but not MHC-II-restricted antigens, we further hypothesize that DCmPs will interact with OVA-specific CD8+ T cells, but not with OVA-specific CD4+ T cells.

To test the hypotheses, we prepared DCmPs from the three approaches and nonstim DCmPs using CFSE-labeled membrane proteins and RhoB-labeled PLGA NPs to study particle binding by T cells. To examine antigen-specific interaction, we incubated bare NPs or coated particles with either B3Z T cells, a murine CD8+ T cell hybridoma with transgenic TCR specific for SIINFEKL/MHC-I molecules,^39,40^ or DOBW cells, a murine CD4+ T cell hybridoma with transgenic TCR specific for OVA_323–339_/MHC-II molecules.^40,41^ (Figure 6A) After 4 hour incubation with fluorescently labeled DCmPs, about 15% B3Z cells are positive for both CFSE and RhoB regardless of the coating approaches. But there were very low frequency of double positive populations found in B3Z cells incubated with nonstim DCmPs, indicating that membrane proteins mediate the binding between DCmPs and B3Z cells.(Figure 6B, 6D) Interestingly, although bare NPs did not show any significant binding to B3Z cells, about 58-66% of B3Z cells are positive for RhoB alone after incubation with DCmPs, significantly higher than the 10% RhoB+ populations observed in nonstim DCmP group.(Figure 6C, 6D) Conversely, only around 1% of DOBW CD4+ cells are found to be double positive after incubation with DCmPs, and neither nonstim DCmPs nor bare NPs treatment showed detectable level of binding.(Figure 6E, 6G) Similar to the observation in B3Z cells, treatment with DCmPs also leads to a noticeable level of RhoB+ DOBW cells, even though DOBW cells did not bind to bare NPs or nonstim DCmPs. (Figure 6F, 6G) Direct comparisons between B3Z cells and DOBW cells regarding the frequencies of double positive populations and RhoB+ populations underscores the preferential binding between DCmPs and SIINFEKL/MHC-I-specific B3Z cells. This highlights that the binding interactions between DCmPs and T cells are primarily mediated by the cognate interaction between MHC and TCRs in our experimental setting. (Figure 6H, 6I)

**Figure 6.**
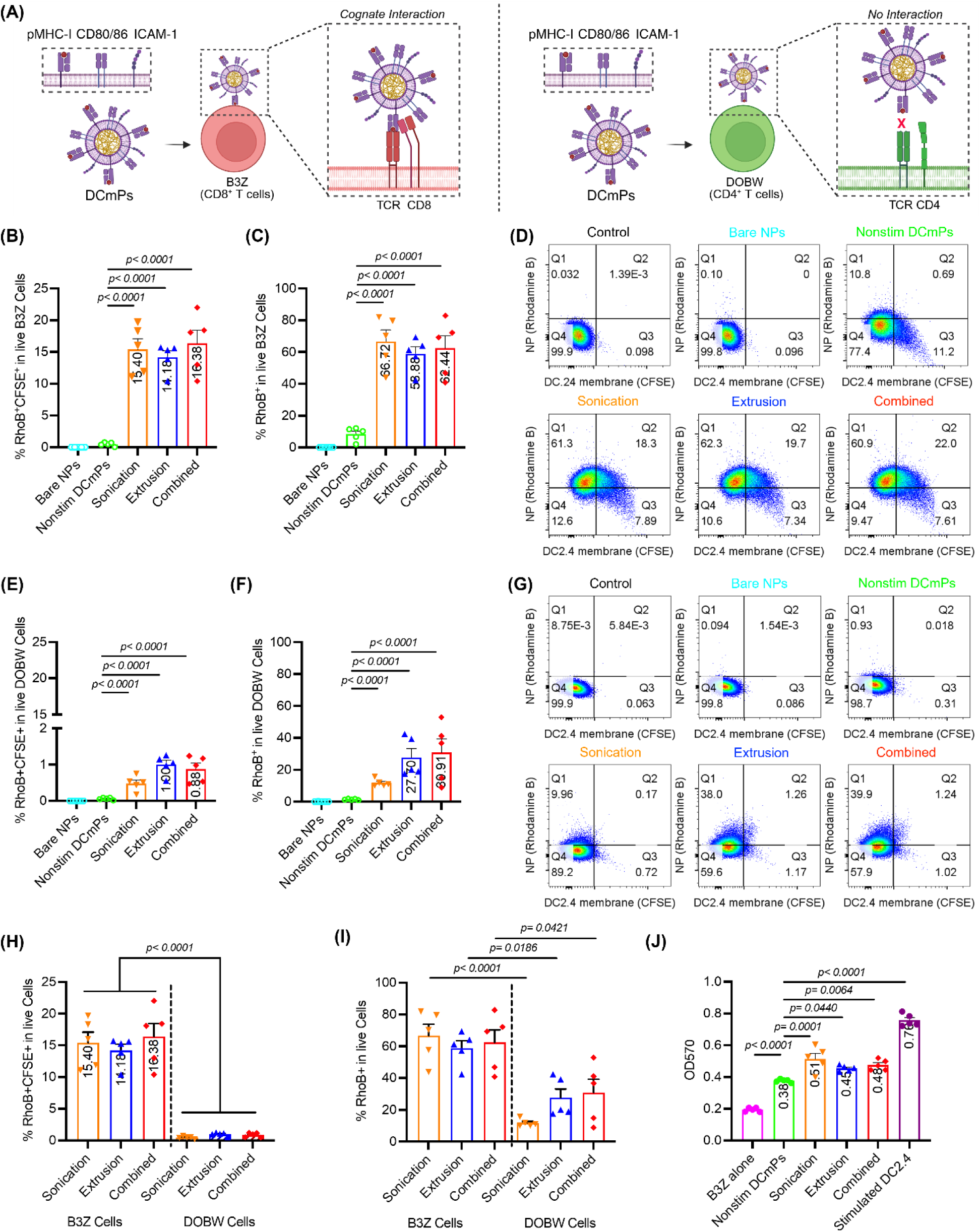
DCmPs preferentially engage antigen-specific B3Z CD8+ T cells and activate B3Z cells via antigen presentation. Fluorescently labeled DCmPs from the three coating approaches were incubated for 4 hours with either B3Z CD8+ T cells or DOBW CD4+ T cells, and the binding between DCmPs and T cells was assessed via flow cytometry. **(A)** Schematic illustration of the T cell binding assay. After 4-hour incubation with DCmPs, the frequencies of B3Z CD8+ T cells that positive for **(B)** both RhoB (PLGA NPs) and CFSE (membrane proteins) or **(C)** RhoB (PLGA NPs) are determined via flow cytometry; **(D)** Representative flow plots for each group are shown. The frequencies of DOBW CD4+ T cells are positive for **(E)** both RhoB (PLGA NPs) and **(F)** CFSE (membrane proteins) are determined via flow cytometry after 4-hour incubation with DCmPs. **(G)** Representative flow plots are shown. Comparison between B3Z and DOBW cells based on: **(H)** double positive (RhoB+/CFSE+) populations; **(I)** RhoB-positive populations. Graphs are plotted based on Panel B, C, E, and F. **(J)** The activation levels of B3Z T cells when stimulated by DCmPs, nonstim DCmPs, and stimulated DC2.4 cells after 48-hour incubation. N=5 for all experiments. Statistical significance was determined using one-way ANOVA followed by Tukey’s multiple comparison test.

SIINFEKL/MHC-I presentation to B3Z cells will lead to express β-galactosidase, which is under the control of interleukin 2 (IL-2) promoter.^39,40^ Therefore, we further assessed the antigen presentation function of DCmPs using B3Z cells.(Figure 6J) To this end, we measured the expression of β-galactosidase by B3Z cells after 48 hours incubation with DCmPs or coculture with DC2.4 cells. As expected, coculture with OVA/LPS-stimulated DC2.4 cells activated B3Z cells, as evidenced by the high level of β-galactosidase secretion. DCmPs also leads to an elevated level of β-galactosidase secretion by B3Z cells, albeit to a lower extent than DC2.4 cells. The three coating processes did not show significant differences in B3Z activation effect, which is consistent with their similar binding ability to B3Z cells. Interestingly, nonstim DCmPs also induced low but measurable level of B3Z cell activation. We suspect this is caused by the low level of SIINFEKL/MHC-I expression found in non-stimulated DC2.4 cells (Figure 5A) Indeed, we confirmed that non-stimulated DC2.4 cells can also activate B3Z cells. (Figure S13) Therefore, DC2.4 cells may have an innate ability to induce low level of B3Z cell activation, which likely contributed to the B3Z activation effect by nonstim DCmPs. Since DC2.4 cells can not present MHC-II molecules and DCmPs don’t bind to DOBW cells to a significant level, (Figure 6E, 6F, 6G) we also confirmed that, OVA/LPS-stimulated DC2.4 cells cannot activate OVA_323–339_/MHC-II-specific DOBW cells unlike BMDCs. (Figure S14) In conclusion, we demonstrate that DCmPs acquire the MHC protein functions and preferentially bind to antigen-specific T cells and facilitate antigen presentations.

## 4. Discussion

Membrane-coated particles excel at cell-mimicking interfacing ability for they acquire source cell membrane protein functions.^1–4^ DC membrane-coated particles, in particular, can carry many important DC membrane proteins, including MHC molecules, co-stimulatory proteins (CD80/86) and adhesive molecules (CCR-7/ICAM-1), which allows DCmPs to engage and activate T cells for antigen-specific therapy similar to DCs.^8,21–23^ Unlike dendritic cell-based therapy,^42^ DCmPs are potentially more flexible for various routes of administration to facilitate tissue-specific targeting and allow membrane protein modification to incorporate *de novo* functions.^17–20^ Owing to these unique advantages, DCmPs have recently been explored for T-cell modulations. ^8,21–23^ To further advance this technology for antigen-specific therapeutics, it is essential to produce coated particles with high consistency in size distribution, the coated protein amount, and the composition of DCmP product. However, our current understanding of the membrane coating process remains limited, which necessitates the experimental approach to identify the optimal coating process.^3,4,32–34^ Moreover, partly constrained by the available technologies, available quantitative analyses of the coating coverage and the amount of unbound protein contents in the final products have been sparse.^5,7^ The direct comparison between sonication and extrusion coating approaches has also been limited.

To address this gap in our knowledge, and to optimize the coating process for DCmPs, we coated PLGA NPs with DC2.4 cell membrane proteins using both sonication and extrusion processes. Additionally, we developed a combined coating approach during which the sonicated particle suspension was subsequently extruded after centrifuge purification. We hypothesize that this combined approach can improve particle size uniformity while maintain high amount of coated proteins. Indeed, the combined approach yielded comparatively lowest diameter increase and the narrowest size distribution among the three coating methods, suggesting that combined approach potentially exerts better control over the size of membrane-coated nanoparticles. (Figure 2B, 2C) Additionally, although the combined approach produced the smallest coated particles, the combined coating approach still achieved slightly higher amount protein coating (10.9% of feeding protein) relative to extrusion alone(about 5.5% of feeding proteins), with sonication achieving highest amount of protein coating (15.2% of feeding protein).(Figure 3A) Taken together, the combined coating approaches show superior control over the size of DCmPs among the three coating approaches and improve the amount of protein coating relative to extrusion.

To gain a better insight of the composition variation of DCmPs from the three coating approaches, we leveraged the well-established homotypic interaction between membrane-coated particles and source cells and utilized DC2.4 cells to detect the presence of both coated particles and unbound proteins, to circumvent the resolution limitation of flow cytometry and confocal imaging. (Figure 4) Using DCmPs composite of RhoB-labeled PLGA NP and CFSE-labeled membrane proteins, we found that about 77-80% of DC2.4 encapsulated both PLGA cores and membrane proteins after 4h incubation with DCmPs. (Figure 4A, 4D) We conclude that this double positive population likely have internalized membrane-coated particles for two main reasons. Firstly, bare NPs are not readily taken up by DC2.4 cells as shown in Figure 4C. Secondly, when treated with a mixture of NPs and membrane proteins, only about 1% of DC2.4 cells are double positive, far lower than the frequency in DCmP-treated DC2.4 cells. (Figure S10) Besides the double positive population, 12-14% of DC2.4 internalized PLGA NPs and 4-9% of DC2.4 only took up membrane proteins, indicating the presence of unbound proteins and bare particles. (Figure 4B, 4C) Our confocal microscopy corroborates with flow cytometry experiments and shows that 60-79% of RhoB-labeled PLGA NPs colocalize with CFSE-labeled membrane proteins, confirming that majority of PLGA nanoparticles within the final products are coated with membrane proteins. (Figure 4E, 4F) Additionally, about 50% of CFSE-labeled proteins are unbound in samples from all three coating approaches according to confocal imaging. (Figure 4E and 4G) The discrepancy in the composition of DCmP products determined using flow cytometry and confocal microscopy can be caused by several factors. Firstly, the relatively low sample numbers in confocal imaging can lead to sampling bias and reduce the accuracy in determining the composition. Secondly, the homotypic interaction promotes membrane protein internalization by DC2.4, thus lowering the possibility of identifying protein alone populations using flow cytometry. In contrast, confocal imaging provided a more detailed look of intracellular proteins, thus able to distinguish unbound proteins from coated proteins within cells. Nevertheless, our work represents a new strategy to determine the composition of membrane-coated particles. We show that, for all three coating approaches, about 80% of PLGA particles are membrane-coated NPs and the final products also contain a considerable amount of unbound proteins despite of the centrifugation purification process.

DCmPs can acquire the ability to activate T cells likely through pMHC-mediated antigen presentation for they carry critical presentation-related membrane proteins, including MHC, CD80/86, and ICAM-1. ^8,21–23^ Using OVA proteins as the model antigen, we identified an LPS-stimulation condition that significantly upregulated these key proteins in DC2.4 cells and confirmed the presence of these proteins on DCmPs. (Figure 1, 3, S1) While the sonication and combined approach showed a higher overall quantity of coated proteins, our western gel analysis was not conclusive on how coating approaches influences the coating of individual proteins. (Figure 3C, S8) Further studies are still required for this aspect. But we confirmed that LPS-stimulation is critical in improving the amount of key surface protein coated on DCmP, consistent with another study using BMDC membrane proteins.^22^ Through flow cytometry, we detected significantly higher level of SIINFEKL/MHC-I, overall MHC-I, CD86, and ICAM-1 on DCmPs than on nonstim DCmPs. (Figure 5) Thus, the amount of individual proteins coated on DCmPs can be improved using adjuvant-stimulated DCs, but the impact of coating approaches is not significant in our study.

We confirmed that DC2.4 cells exclusively present MHC-I antigens using multiple assays, including Eα peptide presentation and DOBW activation. (Figure S3, S12, S14) The exclusive MHC-I expression dictates that DC2.4 cells will preferentially bind to antigen-specific CD8+ T cells. We thus assessed the ability of DCmPs to selectively bind to antigen-specific CD8+ T cells and to present antigens to CD8+ T cells, using SIINFEKL/MHC-I-specific CD8+ B3Z T cells and OVA_323–339_/MHC-II-specific CD4+ DOBW T cells as model. After 4 hour incubation with fluorescently-labeled DCmPs, about 15% of CD8+ B3Z cells acquired DCmPs, whereas only 1% of CD4+ DOBW cells are double positive. Nonstim DCmPs showed no detectable binding to either cell lines. (Figure 6B, 6E, 6H) These data strongly suggest that it is the cognate interaction between pMHC-I and TCRs that mediates the preferential association between DCmPs and B3Z cells. Furthermore, DCmP treatment of B3Z cells lead to the production of β-galactosidase, confirming that DCmPs can present SIINFEKL antigens and activate B3Z cells. (Figure 6J) Nonstim DCmPs exhibited detectable but reduced level of B3Z activation compared to DCmPs, likely due to the presentation ability of non-stimulated DC2.4 cells. (Figure 5B, S13) Unexpectedly, about 60% of B3Z cells only showed detectable level of RhoB, suggesting binding to bare PLGA NPs. (Figure 6C) A relatively lower frequency of DOBW cells (about 30%) are also found to be only RhoB positive. (Figure 6F, 6I) This is contradictory to the fact that neither bare NPs nor nonstim DCmPs showed significant binding to either B3Z cells or DOBW cells.

(Figure 6C, 6F) Given that mixing proteins and NPs did not improve bare PLGA NP engagement to cells, (Figure S10) we speculate that the observed RhoB+ populations in B3Z and DOBW cells are caused by the uptake of PLGA NPs that carried low amount of DC membrane proteins. These membrane proteins may mediate interactions between particles and cells, but their quantity could be below the detection threshold of flow cytometry. The accessory molecules, such as CD86 and ICAM-1, may have contributed to the non-antigen-specific binding between particles and DOBW cells. However, testing this hypothesis will require high resolution characterization of DCmPs to confirm the presence of small quantity of coated proteins, which is beyond the scope of current study. Future studies will also be needed to elucidate the role of accessary molecules in T-cell binding. Nevertheless, our data clearly demonstrates that DCmPs preferentially bind to antigen-specific T cells *in vitro* and can activate T cells via antigen presentation.

## 5. Conclusion

We systemically compared three coating approaches, i.e. sonication, extrusion, and the combined approach, for the manufacture of DCmPs using DC2.4 cell membrane proteins, and demonstrated that the combined approach achieved the best control over particle size among the three approaches and improved overall coated protein amount over extrusion alone. Using DC2.4 as a surrogate, we confirmed all three coating approaches achieve similarly high level of protein coating on PLGA NPs and the final products contain unbound proteins and low percentage of uncoated particles. The presence of presentation-related proteins on DCmPs are confirmed using western blot and flow cytometry, and LPS-stimulation improved the amount of coated proteins on DCmPs. Finally, we demonstrate that DCmPs preferentially bind to cognate T cells likely mediated by the cognate MHC/TCR interaction and are capable of antigen presentation to T cells. Our findings establish a new coating strategy and a characterization strategy for the development of membrane-coated particles, and demonstrate the potential of DCmP technology for antigen-specific therapy.

## Acknowledgement

We would like to acknowledge Dr. Abrar Aljiboury at the BioImaging Center of Syracuse University for her support with confocal imaging and co-localization analysis. We thank Dr. Benjamin Zink at the Transmission Electron Microscopy (TEM) Core Facility of Upstate Medical University for the assistance in acquiring TEM images, and Dr. Lisa Phelps at the Flow Cytometry Core Facility of Upstate Medical University for her expert support on flow cytometry. We thank Dr. David Carnes at the Laboratory Animal Facilities of Syracuse University for his support in supplying C57BL/6 mice tissues. We also extend our sincere appreciation to Dr. James Moon at the University of Michigan for kindly providing B3Z cells, and to Dr. Clifford Harding at Case Western Reserve University for the generous gift of DOBW cells. The illustrations were created in BioRender (https://BioRender.com/u08ltfd). This research is supported by the Syracuse University Startup Fund.

## Author Contributions

S. P. and Y. W. conceived the project, designed the experiments and wrote the manuscript. S. P., S. S. J., A. Z. and R. R. B. performed particle synthesis and characterization. S. P. and A. Z. validated DC stimulation. S. P. and J. B. W. performed protein characterization. M. M. cultured BMDCs. B. V. maintained cell line. S. P. and A. M. S. performed membrane protein isolation. Y.W., X. H., and J. L. H. provided resources and supervision. All authors reviewed and approved the manuscript.

## Data Availability Statement

The data that support the findings of this study are available from the corresponding author upon request.

Received: ((will be filled in by the editorial staff))

Revised: ((will be filled in by the editorial staff))

Published online: ((will be filled in by the editorial staff))

