## Supplemental Figures for "Development of Dendritic Cell Membrane-Coated Nanoparticles for Antigen-Specific T Cell Engagement"

**Supplementary Information**

*Sao Puth,[1, 2] Shruti Sunil Jadhav,[1, 2] Ali Zareein,[1, 2] Jimmy Blauser-Wilson,[1, 2] Mina Mahmoudi,[1, 2] Ruben Rojas Betanzos,[1,2] Bayonel Ventura,[1,2] Andrea M. Sprague-Getsy,[2,3] Xiaoran Hu,[2,3] James L. Hougland,[2,3,4] Yaoying Wu[1, 2, 5]**

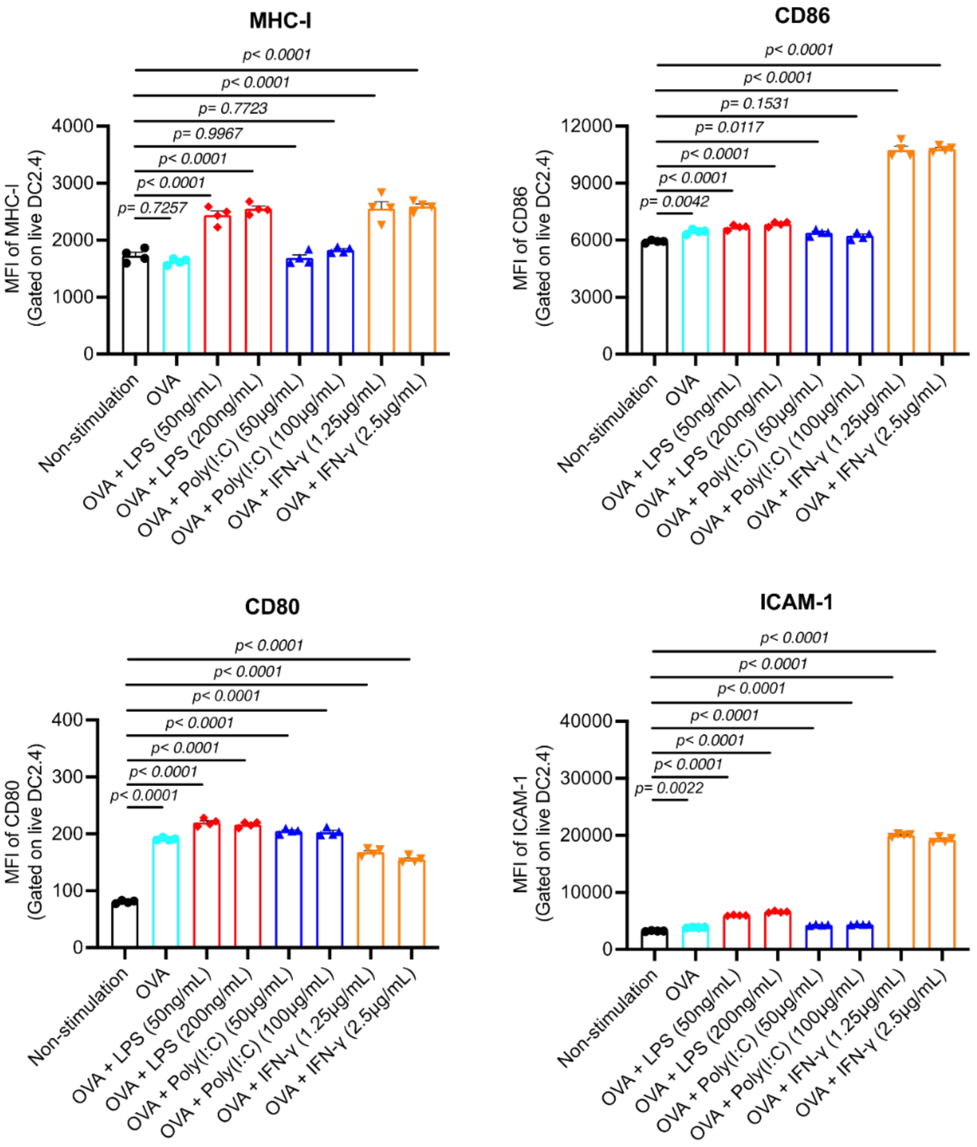


**Figure S1. The expression level of key presentation–related membrane proteins in DC2.4 cells after the stimulations of LPS, Poly(I:C), and IFN-γ.** DC2.4 cells were stimulated with OVA protein (0.3 mg/mL) in combination with various adjuvants at different concentrations: LPS [50 or 200 ng/mL], Poly(I:C) [50 or 100 μg/mL], and IFN-γ [1.25 or 2.5 μg/mL] for 15 hours. The expression level of proteins, including MHC-I, CD80, CD86, and ICAM-1, were assessed using flow cytometry. N=4. Statistical significances were determined using one-way ANOVA followed by Tukey's multiple comparison test.

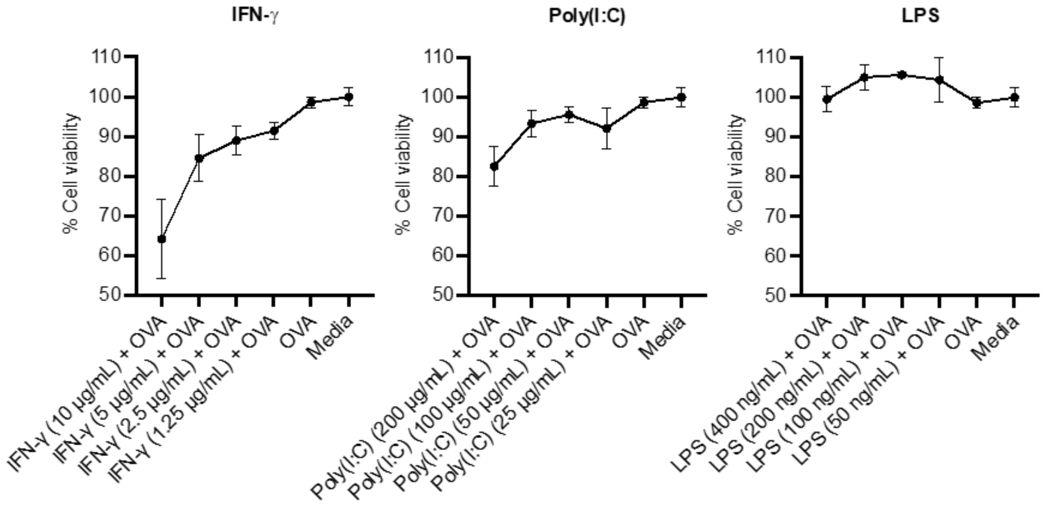


**Figure S2. Cytotoxicity of adjuvant stimulation in DC2.4 cells.** DC2.4 cells were stimulated with OVA protein (0.3 mg/mL) in combination with various adjuvants at different concentrations: IFN-γ [1.25, 2.5, 5, or 10 μg/mL], Poly(I:C) [25, 50 100, or 200 μg/mL], and LPS [50, 100, 200, or 400 ng/mL] for overnight. Cytotoxicity was assessed using the MTS assay. The absorbance at 490 nm was determined using a plate reader. Cell viability was calculated and expressed as a percentage using the formula: ($\frac{sample}{live\ cell\ control} \times 100\%$).

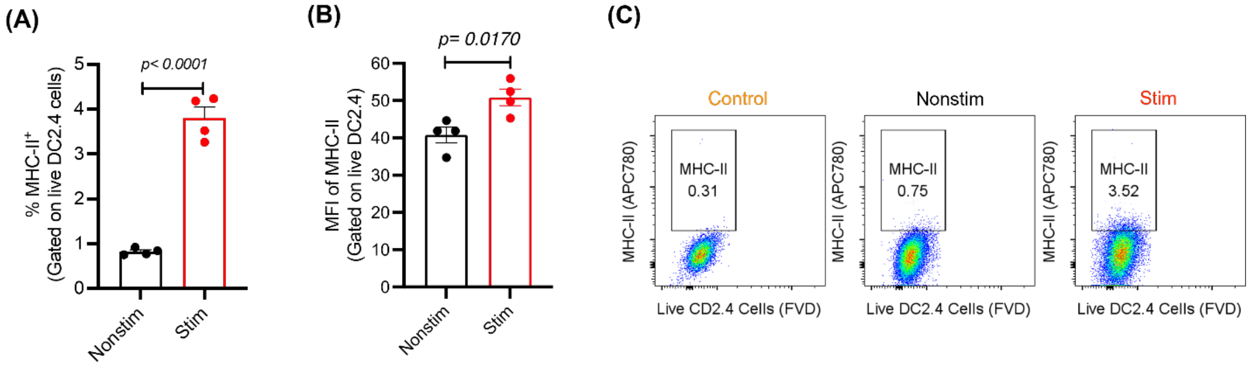


**Figure S3. Comparison of MHC-II expression in non-stimulated and stimulated DC2.4 cells.** The expression of MHC-II on non-stimulated (Nonstim) and stimulated (Stim) DC2.4 cells was determined using flow cytometry. Quantification of MHC-II expression: **(A)** percentage of MHC-II+ cells and **(B)** mean fluorescence intensity (MFI) of MHC-II expression. **(C)** The representative flow cytometry plots. N=4. Statistical significance was determined using Student's *t*-test.

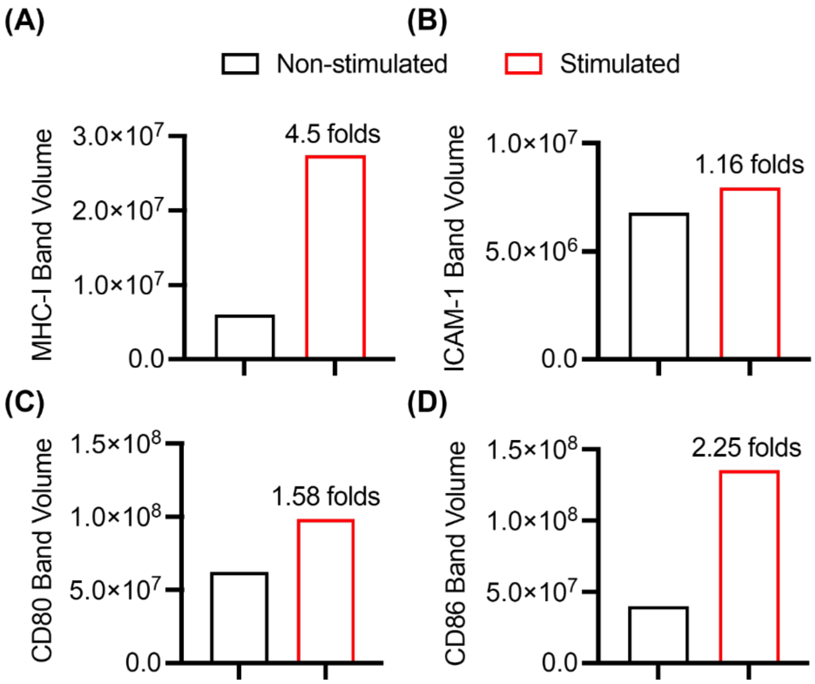


**Figure S4: Western blot band volume of surface membrane proteins in DC2.4 cell lysates.** Quantitative analysis of western blot band intensities based on Figure 1C was determined using ImageJ software. Protein levels of **(A)** MHC-I, **(B)** ICAM-1, **(C)** CD80, **(D)** CD86 were normalized to $Na^+/K^+$-ATPase and the fold increases relative to the non-stimulated control are shown.

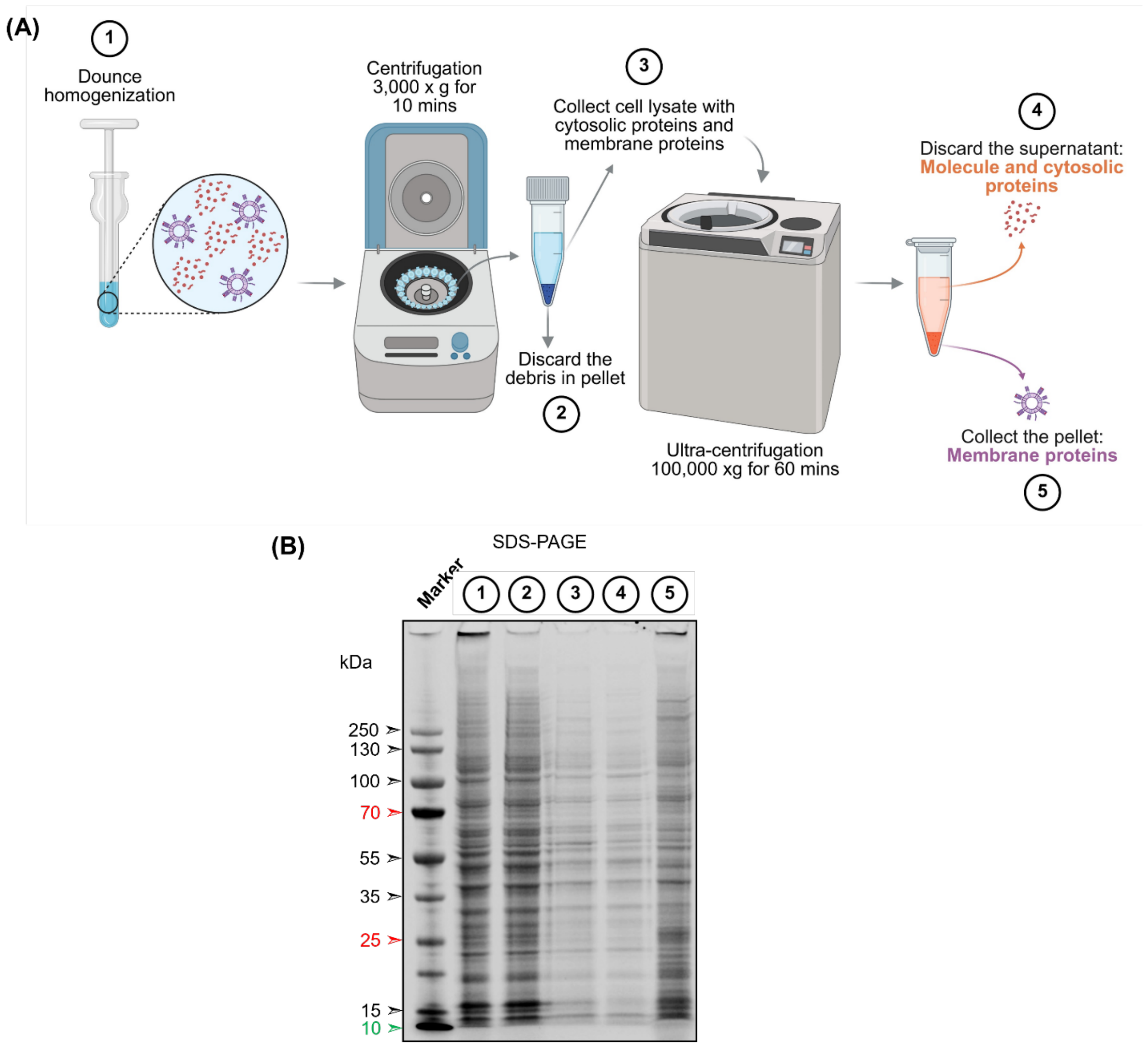


**Figure S5. Workflow and validation of DC2.4 membrane protein extraction.** Membrane proteins from DC2.4 cells were isolated using Dounce homogenization followed by ultracentrifugation. **(A)** Schematic illustration of the DC2.4 membrane isolation workflow. **(B)** SDS-PAGE gel images of the protein profile at each step of the extraction process.

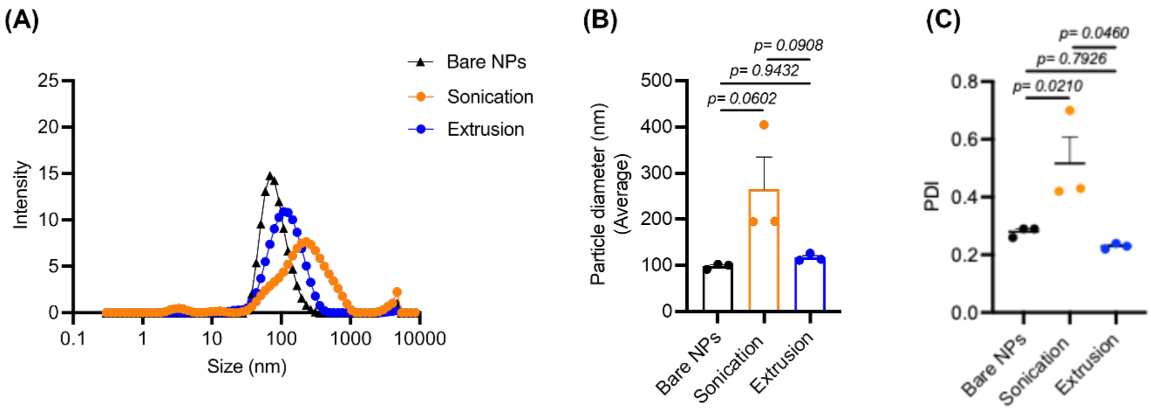


**Figure S6. DLS analysis of DC2.4 membrane-coated PLGA nanoparticles prior to centrifugation.** Membranes were coated onto PLGA nanoparticles using either sonication or extrusion at a 1:1 weight ratio (w/w). Without removing the unbound proteins by centrifugation, the obtained particle suspensions were characterized using DLS. DLS analysis showing **(A)** particle size distribution, **(B)** average particle size, and **(C)** polydispersity index (PDI). N=3. Statistical significance was determined using one-way ANOVA followed by Tukey's multiple comparison test.

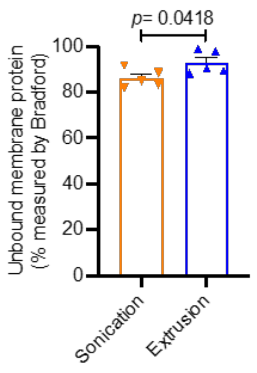


**Figure S7. Comparison of unbound membrane protein in the supernatant after centrifugation of DCmPs prepared by sonication or extrusion.** The amount of unbound membrane protein in the supernatant after centrifugation was quantified by Bradford assay. N=5. Statistical significance was determined using Student's *t*-test.

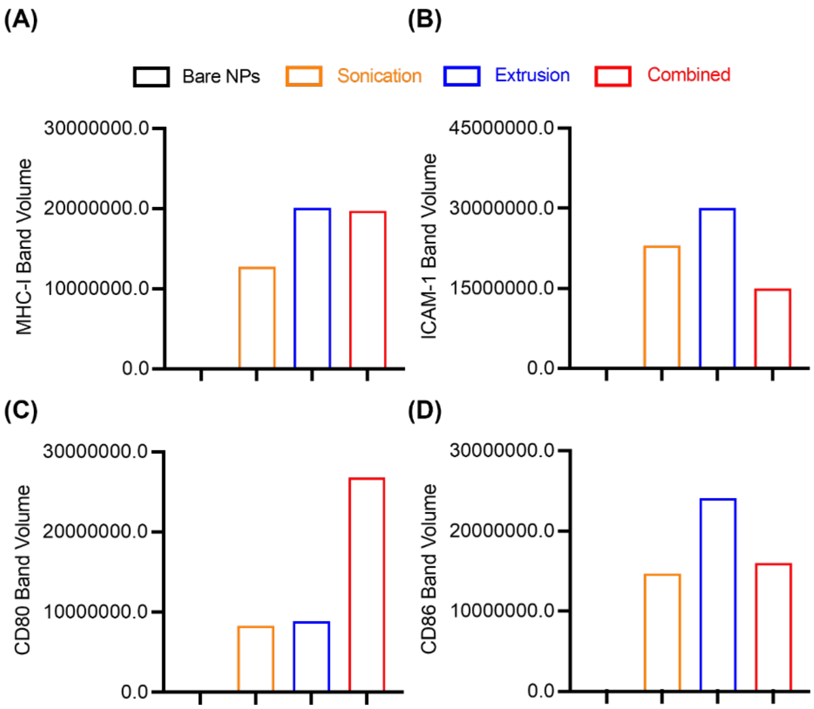


**Figure S8. Quantification of surface membrane proteins on DCmPs.** Quantitative analysis of Western blot band intensities (Figure 3C) was performed using ImageJ software. Expression levels of **(A)** MHC-I, **(B)** ICAM-1, **(C)** CD80, **(D)** CD86 were normalized to $Na^+/K^+$-ATPase as a loading control.

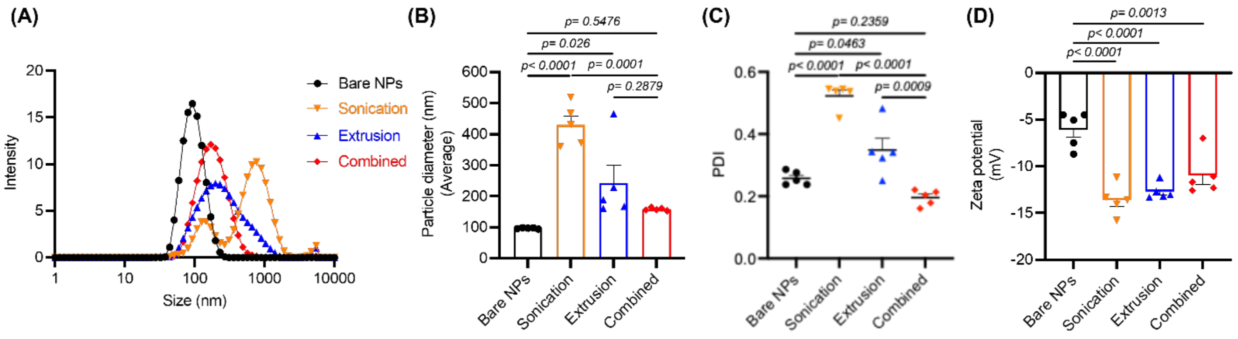


**Figure S9. Characterization of DCmPs composed of RhoB-loaded PLGA NPs and CFSE-labeled membrane proteins.** CFSE-labeled membranes isolated from stimulated DC2.4 cells were coated onto Rhodamine B (RhoB)-labeled PLGA NPs using sonication, extrusion, or a combined process to produce DCmPs. Dynamic light scattering (DLS) analyses were conducted to assess: **(A)** particle distribution, **(B)** average particle size, **(C)** polydispersity index (PDI), and **(D)** zeta potential of both bare NPs and DCmPs. N=5. Statistical significance was determined using one-way ANOVA followed by Tukey's multiple comparison test.

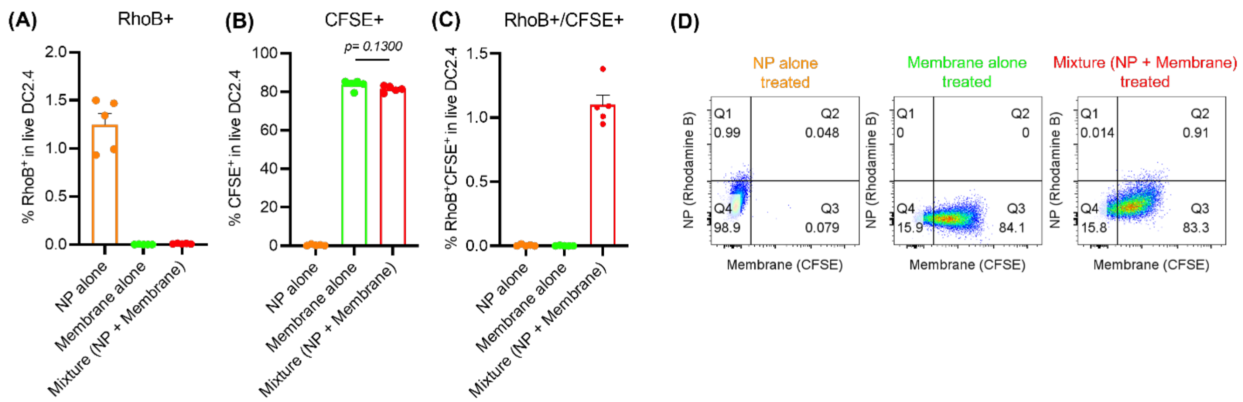


**Figure S10. Binding of nanoparticles, membrane, or membrane-nanoparticle mixtures by DC2.4 cells.** CFSE-labeled membranes isolated from stimulated DC2.4 cells, Rhodamine B (RhoB)-labeled PLGA NPs, or a mixture of both were incubated with DC2.4 cells for 4 hours. Flow cytometry was performed to quantify the percentage of DC2.4 cells that are **(A)** RhoB-positive, **(B)** CFSE-positive, **(C)** double-positive ($RhoB^+/CFSE^+$) populations. **(D)** Representative flow cytometry plots are shown. N=5. P value was determined using one-way ANOVA.

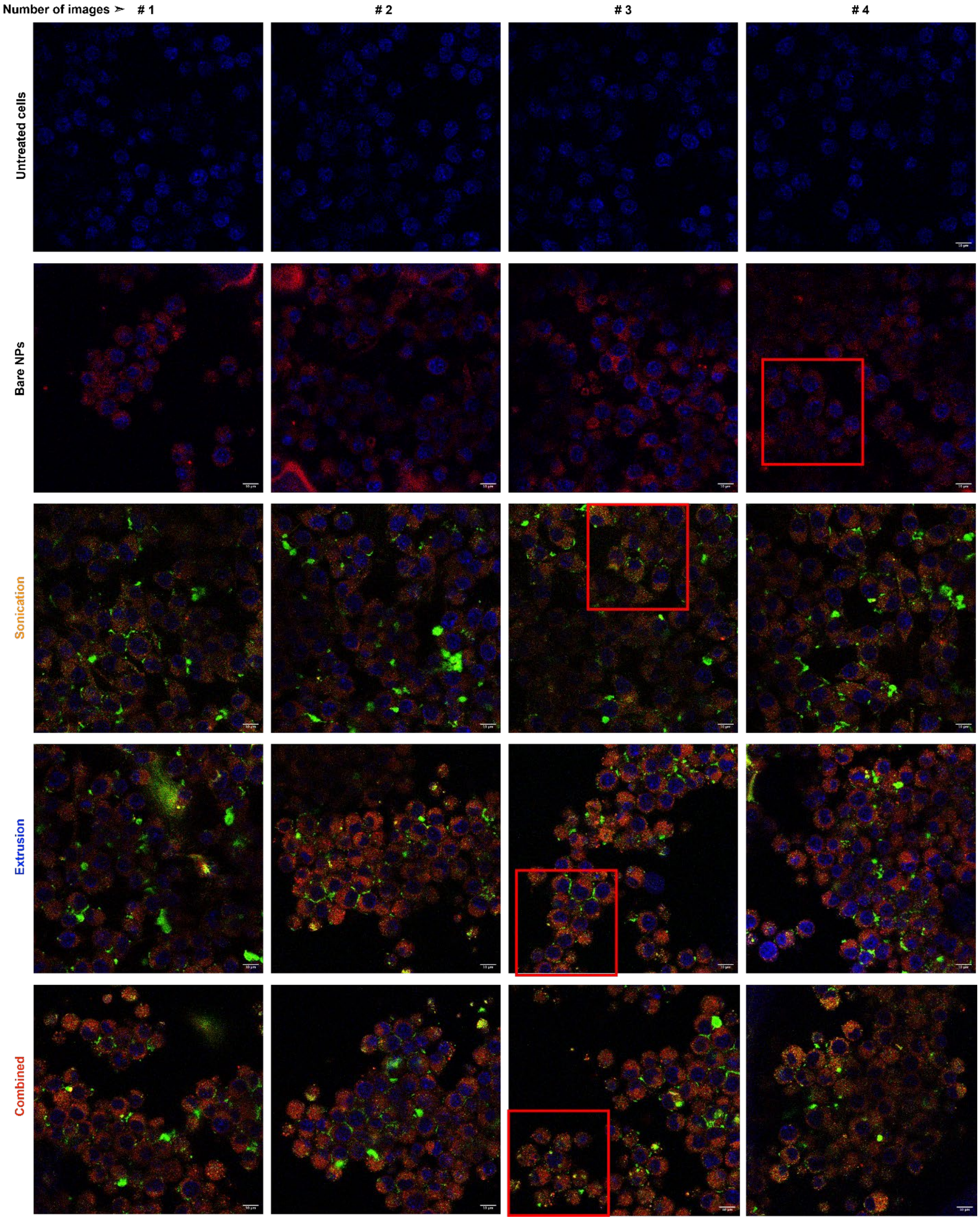
Number of images ➢ # 1
# 2
# 3
# 4
Untreated cells
Bare NPs
Sonication
Extrusion
Combined

**Figure S11. Confocal laser scanning microscopy (CLSM) analysis of DCmPs internalization by DC2.4 cells.** Fluorescently labeled DCmPs were incubated with DC2.4 cells for 4 hours. RhoB-labeled PLGA NPs alone and untreated cells were included as controls. Co-localization of CFSE-labeled membrane proteins (green) with RhoB-labeled PLGA cores (red) was assessed using CLSM and analyzed with ImageJ software. Blue represents for nucleus (DAPI). Four independent images were acquired per experimental group, and representative areas (highlighted by squared-red boxes) are shown in Figure 4E.

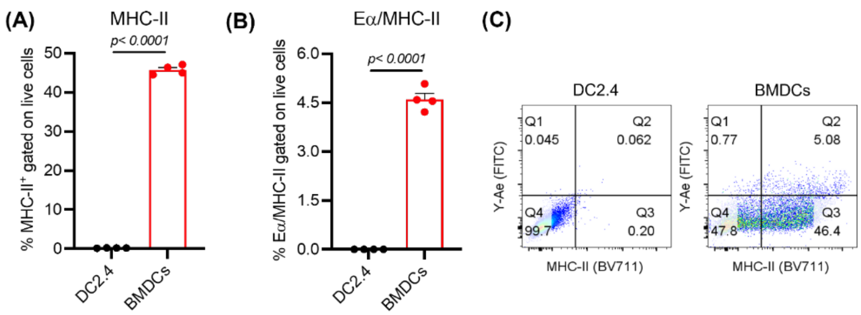


**Figure S12. Comparison of MHC-II and Eα/MHC-II expression between DC2.4 cells and BMDCs.** DC2.4 cells and BMDCs were stimulated overnight with Eα peptide (50 μg/mL) and LPS (50 ng/mL). The expression levels of Eα/MHC-II complexes were assessed using Y-Ae antibodies. All gatings are based on live cells. The percentages of **(A)** MHC-II, **(B)** Eα/MHC-II cells, and **(C)** representative flow cytometry plots, are presented. N=4. Statistical significance was determined using Student's *t*-test.

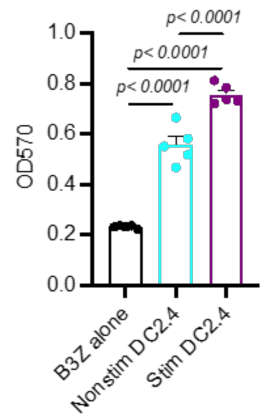


**Figure S13. B3Z activation by stimulated and non-stimulated DC2.4 cells.** B3Z cells ($2 \times 10^5$) were co-cultured with non-stimulated DC2.4 cells (Nonstim DC2.4) or DC2.4 cells stimulated with OVA protein (0.3 mg/mL) and LPS (50 ng/mL) (Stim DC2.4) at 1:1 ratio. After 2 days of co-culture, The activation levels of B3Z T cells were measured using the CPRG/lysis buffer assay. B3Z cell alone was used a control group. N=5. Statistical significance was determined using one-way ANOVA followed by Tukey's multiple comparison test.

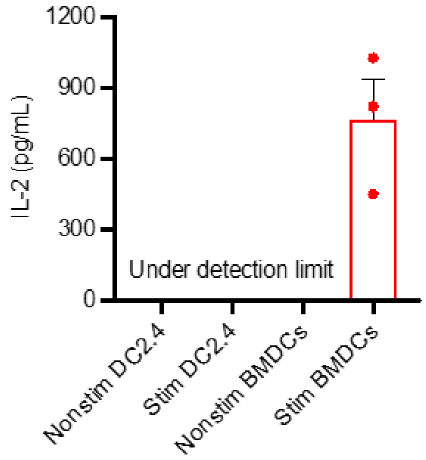


**Figure S14. DOBW cell activation by DC2.4 cells or BMDCs.** DC2.4 cells and BMDCs were stimulated overnight with OVA protein (0.3 mg/mL) and LPS (50 ng/mL) for overnight, before co-cultured with DOBW T cells for 3 days at 1:1 ratio. IL-2 secretion by DOBW cells was considered as an indicator of T-cell activation, and was quantified using IL-2 cytokine ELISA. Non-stimulated DC2.4 and non-stimulated BMDCs served as controls. While BMDCs induced robust IL-2 production by DOBW cells, DC2.4 cells failed to elicit detectable IL-2 response.
